# Computational repurposing of oncology drugs through off-target drug binding interactions from pharmacological databases

**DOI:** 10.1101/2023.07.01.547311

**Authors:** Imogen Walpole, Farzana Y Zaman, Peinan Zhao, Vikki M. Marshall, Frank Lin, David M. Thomas, Mark Shackleton, Albert A. Antolin, Malaka Ameratunga

## Abstract

**PURPOSE:** Systematic repurposing of approved medicine for another indication represents an attractive strategy to accelerating drug development in oncology. Herein we present a strategy of combining biomarker testing with drug repurposing to identify new treatments for patients with advanced cancer.

**METHODS:** Tumours were sequenced with Illumina TruSight Oncology 500 (TSO-500) platform or the FoundationOne® CDx panel. Mutations were manually screened by two medical oncology clinicians and pathogenic mutations were categorised with reference to the literature. Variants of unknown significance were classified as potentially pathogenic if a plausible mechanism and computational prediction of pathogenicity existed. Gain of function mutations were evaluated through the repurposing databases Probe Miner, the Broad Institute Drug Repurposing Hub (Broad Institute DRH) and TOPOGRAPH. Gain of function mutations were classified as repurposing events if they were identified in Probe Miner, were not indexed in TOPOGRAPH which captures active clinical trial biomarkers and excluding mutations for which a known FDA-approved biomarker label exists. The performance of the computational repurposing approach was validated by evaluating its ability to identify known FDA-approved biomarkers. Exploratory functional analyses were performed with gene expression data and CRISPR-dependency data sourced from the DepMap portal. The total repurposable genome was identified by evaluating all possible gene-FDA drug approved combinations in the Probe Miner dataset.

**RESULTS:** The computational repurposing approach was highly accurate at identifying FDA therapies with known biomarkers (94%). Using a real-world dataset of next-generation sequencing molecular reports (n = 94) and excluding the identification of mutations that would render patients eligible for FDA-licensed therapies or local clinical trials, it was found that a meaningful percentage of patients (14%) would have an off-label therapeutic identified through this approach. Exploratory analyses were performed, including the identification of drug-target interactions that have been previously described in the medicinal chemistry literature but are not well known, and the evaluation of the frequency of theoretical drug repurposing events in the TCGA pan-cancer dataset (73% of samples in the cohort).

**CONCLUSION:** Overall, a computational drug repurposing approach may assist in identifying novel repurposing events in cancer patients with advanced tumours and no access to standard therapies. Further validation is needed to confirm the utility of a precision oncology approach using drug repurposing.

## Introduction

Improving access to novel therapeutics for patients with advanced and poor prognosis cancer is important and challenging. Twin challenges in these patients are access to novel therapeutics and limited drug efficacy. The traditional pathway for accessing novel therapeutics in this patient population is through phase I clinical trials. However, several issues limit the utility of this pathway to accelerate drug development in this subgroup of patients, including access, logistics and poor efficacy.

As a consequence of safety being the primary endpoint for these trials, strict inclusion criteria often exclude patients from enrolment, with studies showing that as few as 34% of real-world cancer patients are able to access these trials (1). Access is compounded by lack of locally available clinical trials for up to 77% of patients (2). Logistically, due to cohort size limitations in dose escalation, phase I trials are not continuously accruing patients, further limiting trial availability. Finally, despite nearly doubling of efficacy rates on aggregate across phase I trials over the last 20 years, efficacy remains low with response rates of 12% (3).

To overcome limited efficacy, biomarker enriched clinical trials consistently report higher response rates than biomarker-agnostic studies (4). This approach is consistent with the objective of precision oncology, which aims to identify targetable molecular aberrations that are unique to subsets of patients (5). Consequently, trends in anti-cancer drug development for patients with advanced solid tumours are increasingly aiming to match patients with specific molecular aberrations to trials targeting those aberrations.

To mitigate issues relating to the relative low frequency of these aberrations, trials with master protocols using ‘umbrella’ (single condition, multiple sub-studies for different molecular aberrations) and ‘basket’ (multiple conditions, single targetable molecular aberrations) have been designed to improve this matching process (6). The NCI-MATCH master protocol is one example of a highly successful umbrella study, identifying ‘actionable’ alterations in 38% of patients screened for this study (7).

However, despite the significant cost and infrastructure development incorporated in these master protocols, results to date highlight two specific problems hindering drug development. Firstly, despite 38% of patients having actionable alterations, only 18% of patients were assigned treatment (7) following the application of exclusion criteria. Secondly, although some sub-studies in NCI-MATCH have been an unequivocal success such as *BRAF* targeted therapy in non-gastrointestinal tumours, other therapies have modest response rates as low as 3%, although response in specific tumour subtypes may assist in biomarker refinement (8).

A drug repurposing approach, defined as identifying new cancer indications for existing approved drugs, is an attractive alternative. Specifically, repurposing enables wider access to therapies which are widely available for other indications, with established safety profiles and prescribing guidelines enabling more widespread clinical use of repurposed drugs. Additionally, repurposing, if successful, has numerous well documented financial and logistical advantages compared to traditional drug development (9). Although repurposing has been successful for non-cancer indications, such as sildenafil, which was developed for angina and has been licensed for treatment for erectile dysfunction (10), few licensed repurposed therapies for oncology indications exist, with thalidomide in myeloma being the best-described example (9).

Most repurposing efforts have been developed in response to *in vitro* and *in vivo* preclinical studies demonstrating activity of specific compounds or by serendipity (9, 11). *In vitro* systematic drug repurposing screens have systematically evaluated compounds for anti-cancer activity, with the resource mostly being used for target identification (11). The Broad Repurposing Hub is one such systematic screen, which involved the curation of 4,707 compounds, experimental confirmation of their identities and annotated these compounds with reference to the literature (12). Overall, three broad types of drug repurposing have been described: off-label use for the same molecular aberration in a different indication (such as the repurposing of trastuzumab from human epidermal growth factor receptor (HER2) amplified breast cancer to HER2 amplified gastric cancer); off-target activity (such as the use of imatinib to target *KIT* mutations in gastrointestinal stromal tumours); and combination approaches based on *in vitro* assays (13).

To our knowledge, a systematic evaluation of off-target activity of FDA-approved drugs to target molecular variants for which approved therapies do not exist has not been undertaken. We have previously developed a computational, objective, quantitative assessment of small molecules for their use to selectively study specific proteins, also termed chemical probes, that we named Probe Miner (15). Probe Miner is based on six different quantitative scores of which potency and selectivity have the highest weight in the final prioritization. To calculate the scores, Probe Miner uses publicly available pharmacological data of several resources integrated in the knowledgebase canSAR, that also integrates its own curated pharmacological data from selected publications (16). Accordingly, Probe Miner can identify the most potent and selective compound to study a specific protein. The Broad Institute DRH is an annotated repurposing library combining publicly available clinical-drug structures from regulatory data and public databases with extensive manual curation (12). TOPOGRAPH is a compendium of approved and experimental therapies assembled from regulatory data, public databases and literature review used to meet the clinical need for tiered assessment for actionability and linking biomarkers to clinical trials (17). The Cancer Dependency Map Project builds upon the original Cancer Cell Line Encyclopedia (18), which involved the systematic molecular profiling of 1,000 cell lines, and performed large-scale functional genomics profiling via RNA-interference and CRISPR screens to identify genetic dependencies. In parallel, these cell lines had systematic drug sensitivity profiling performed. Here, we hypothesise that using these publicly available resources to systematically evaluate off-target drug repurposing activity is feasible and potentially expands therapeutic options available to patients.

## Methods

### Patients

Patients with metastatic or advanced solid organ malignancies were referred from participating partner hospitals via the Monash Partners Comprehensive Cancer Consortium (as coordinating body) who were sequenced using next generation sequencing (NGS) through the Molecular Screening and Therapeutics (MoST) clinical trial platform study during patient care were accessed. Ethics approval for the current study was granted through the Alfred Hospital Human Research Ethics Committee (HREC) (419/21).

The MoST trial NGS panel utilises either the Illumina TruSight Oncology 500 (TSO-500) panel or the FoundationOne® CDx panel. The TSO-500 analyses 523 genes for single nucleotide variations (SNVs) and insertion/deletions. It also analyses 55 genes for fusion transcripts and splice variants. Alternatively, the FoundationOne® CDx panel analyses 324 genes for substitutions and insertion/deletions.

Microsatellite instability (MSI) status was classified as high and low as per manufacturer’s specification of the respective panel. Tumour mutational burden (TMB), defined as number of non-synonymous mutations per megabase, was dichotomised at the threshold of 10mut/mb consistent with KEYNOTE-158 study that form the basis of pembrolizumab approval (19).

### Classification of Variants

NGS reports were secondarily reviewed by two medical oncology clinicians (FZ, IW, MA) to classify and curate genomic variants of significance. Variants were classified into either potential gain of function or loss of function mutations based on literature review and annotated as pathogenic, likely pathogenic, or likely benign by cross-referencing against the Catalogue of Somatic Mutations in Cancer (COSMIC) database (20). Their corresponding pathogenicity score by Functional Analysis Through Hidden Markov Models (FATHMM) was recorded (21). A literature review to identify pathogenic mutations which had been previously orthogonally functionally validated was then performed. For variants of uncertain significance (VUS), mutations were annotated as possibly pathogenic if they demonstrated a plausible mechanism, pathogenic FATHMM score, in combination with review by an oncologist (MA) and medicinal chemist (AA). In general, truncating mutations were classified as loss of function and amplifications were classified as gain of function. Non-synonymous single nucleotide (nsSNV) variants were classified on a case-by-case basis. Kinase and hotspot mutations in known oncogenes were classified as potential gain of function mutations.

### Identification of Drug Targets

Gain of function mutations were assessed as possible drug targets using three well-established and accessible complementary databases– Probe Miner (15), the Broad Institute DRH (12) and TOPOGRAPH (17). Probe Miner uses a systematic quantitative approach combining publicly available chemical and pharmacological data with an objective scoring metric, weighting selectivity and potency, to identify chemical probes, including licensed medicines, that bind specific protein targets (15). To assess the predictive performance of Probe Miner, we postulated that the top scoring results from this platform could re-identify FDA-approved drugs (15), which would be heavily weighted for selectivity and potency for the specific protein. FDA approved drugs targeting that mutation were reviewed. A list of drugs meeting these criteria was curated for each gain of function mutation. Gain of function mutations were then filtered using the Broad Institute DRH (12) and TOPOGRAPH databases (17). Again, a list of drugs identified as potentially targeting these gain of function mutations was collated. The Broad Institute DRH was utilised to assess the degree of overlap in repurposing opportunities identified by Probeminer. Variants in TOPOGRAPH were assumed to have preclinical/clinical rationale given this database specifically curates this information.

### Categorisation of Drug Repurposing Events

From the list of curated drugs identified via Probe Miner and the Broad Institute DRH, drug repurposing events were categorised. A *repurposing event* was classified as any gene aberration with a drug repurposing opportunity identified on one of the databases. Initially gain of function mutations in genes with well recognised, investigated and approved targeted therapies (Tier I mutations) were removed after review by two oncologists (IW, MA) and cross-referencing with the Table of Pharmacogenomic Biomarkers in Drug Labelling published by the FDA (22). These included *KRAS*, *ERBB2*, *EGFR*, *BRAF*, *KIT*, *CDK4*, *CDK6* and *PIK3CA*. Mutations of potential clinical significance (aligning with Tier II (23) mutations), for which an active ongoing clinical research program was investigating trial therapies with a strong preclinical/clinical rationale, were removed using TOPOGRAPH (17) and annotated as *repurposing events with trial level evidence*. Remaining mutations were classified as *repurposing events without trial level evidence*. Manual curation of remaining mutations was performed by an oncologist (MA) and a medicinal chemist (AA). This included reviewing mechanistic data to ensure the binding identified in public databases was likely to be functionally consequential. Mutations with identified drugs meeting these criteria were considered novel drug repurposing events from genomic variants. Well-known off-label drug-target interactions were annotated as *off-label repurposing* events (MA, AA) and drug-target interactions which were not well known were annotated as *novel repurposing events*. Analysis was then undertaken to assess the degree of overlap between recommendations generated from all three databases: Probe Miner, Broad Institute and TOPOGRAPH.

### Exploratory functional analysis of drug-target interactions

Exploratory functional analysis was performed using data obtained from the Cancer Dependency Map. Drug sensitivity data from the PRISM repurposing project (11) and genomics of drug sensitivity screen (24, 25) were cross-referenced against gene expression data (26) and CRISPR-gene dependency data (27). Drug-target combinations identified by Probe Miner, for which a possible relationship could be observed, were annotated.

### Evaluation of repurposable genome

The repurposable genome was defined as genes for which a drug repurposing event could be identified by Probe Miner. This was assessed first by converting the Uniprot accession indexed in Probe Miner to an entrez ID and gene name. All compounds indexed in Probe Miner per gene, with a Probe Miner global score >0.25, were cross referenced against FDA licensed therapeutics from the Drugs@FDA database, downloaded on 6 May 2022. The top ranked unique compounds were then collated with corresponding genes, to create the total repurposable genome. The genes included in the total repurposable genome were then evaluated for gene mutations in the The Cancer Genome Atlas (TCGA) pan-cancer analysis of whole genomes dataset (28) from Cbioportal (29, 30) to calculate repurposable frequency. All analysis was performed in R version 4.2.2.

## Results

### Validation of Probe Miner accuracy for FDA approved medications

Drugs that are FDA licenced with a biomarker indication in oncology were curated from the Table of Pharmacogenomic Biomarkers in Drug Labelling published by the FDA (22). This table was manually curated (FZ, MA) to ensure that the drug target for the FDA approved drug was correct. Specific examples of curation include, FDA labels identifying a molecular target related to the tumour type (such as the oestrogen receptor, *ESR1* gene in breast cancer) as opposed to the molecular target (such as CDK4 inhibitors). The curated table is shown in Supplementary Table I, with the molecular targets labelled “Target”. For each drug-target combination, manual curation was performed to find small molecular inhibitors that bind the specific molecular target (n = 67, Supplementary Table II). Of the 67 drug-target combinations, Probe Miner identified 94% as targets, with a quantitative score ranging from 0.19 (ivosidenib for *IDH1*) to 0.80 (ponatinib for *ABL1*) (Supplementary Table II). Thirty drugs were ranked in the top ten chemical probes (45%), highlighting the good performance of the *in silico* approach. It must be stressed that Probe Miner includes many chemical probes that are not licensed drugs as its primary use is in chemical biology, so the performance for this alternative use is significant. Highly selective small molecule inhibitors consistently ranked higher than less selective inhibitors (e.g. osimertinib ranked higher than erlotinib for *EGFR*). Of the four drug-target combination missed by Probe Miner, one interaction was the rarely used toremifene as a selective oestrogen receptor modulator (*ESR1*). The full list of FDA licensed drugs with oncology indications and their Probe Miner score and rank is listed in Supplementary Table II.

Quantification of the performance of Probe Miner for detecting true drug-target interactions was performed. For this analysis, a Probe Miner quantitative score of 0.25 was used as it was slightly greater than the lower bound of the range of global scores of FDA licensed therapies with known biomarkers. For the gold standard, curated FDA-approved drug-biomarker combinations were considered true positives and all non-FDA approved drug-biomarker combinations were considered true negatives. Only compounds indexed in both the Drugs@FDA database and Probe Miner were considered in this analysis because Probe Miner may not contain all the latest FDA-approved drugs, that sometimes take a few months to appear in public pharmacological databases. Only biomarkers for which there were more than one licensed therapy were considered as these were most likely to have sufficient data to inform analysis. . Overall, Probe Miner identified FDA-licensed biomarker-drug combinations with moderate-to-high sensitivity (range 0.5-1) and high specificity (range 0.99-1.00) (Table I). Although the precision of Probe Miner was demonstrably lower, this may be due to the high threshold used for true positives in this analysis (i.e. if a drug-target interaction identified by Probe Miner is clinically efficacious but is not FDA listed, it would be identified as a false positive).

**Table I:**
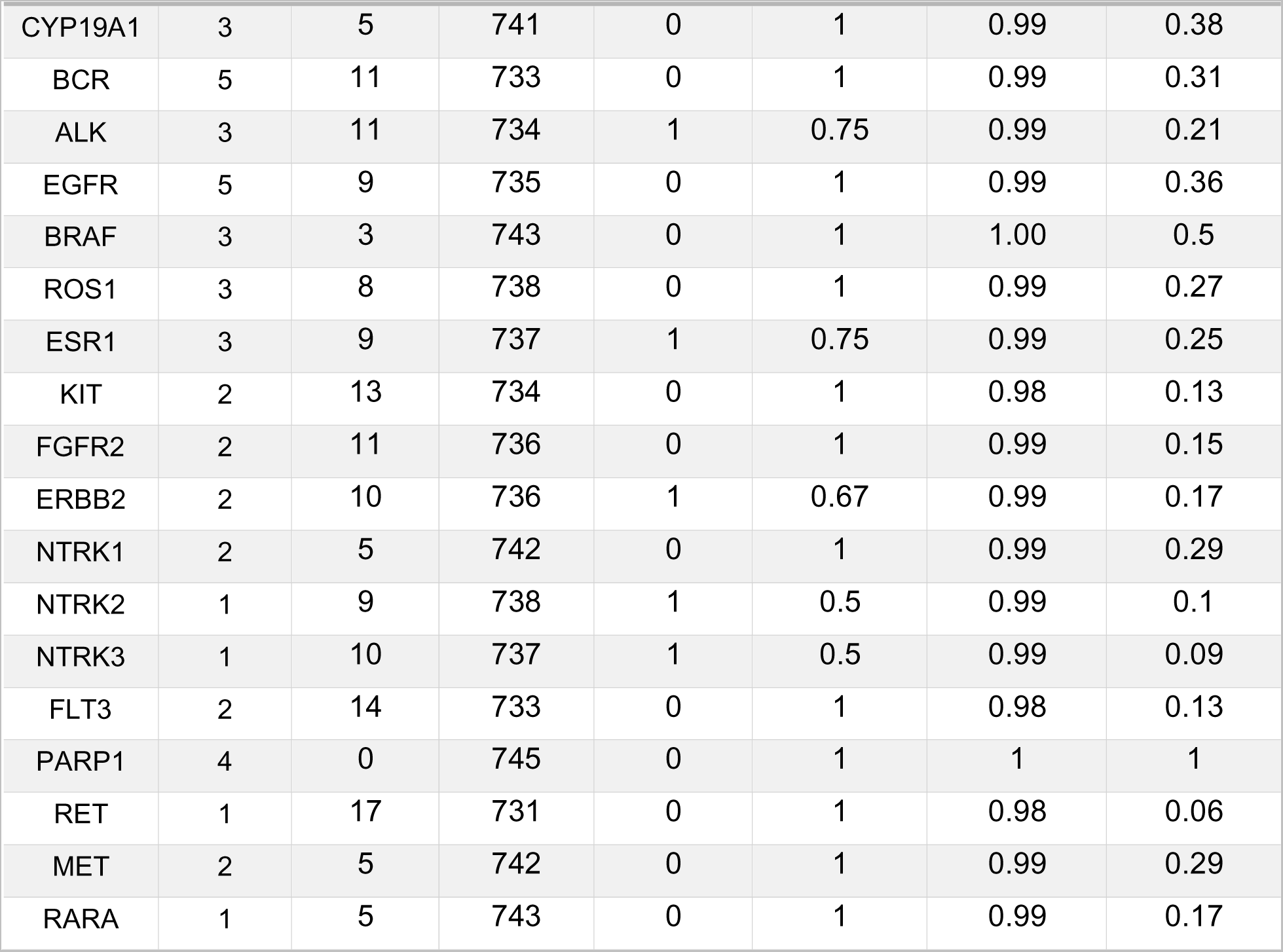
Performance of Probe Miner for identification of FDA-licensed drug-biomarker combinations.

### Patient demographics

A total of 94 patient NGS reports generated from September 2019 to May 2021 were reviewed. The median age of patients undergoing NGS was 63 years with only 36.2% of patients 65 years or older (Table II). The most common tumour group undergoing testing was hepatobiliary/pancreatic cancers (23.4%), followed by gynaecological (21.3%), lower gastrointestinal (12.8%) and upper gastrointestinal cancers (7.4%) (Table II). The 15 subtypes listed under other included carcinoma of unknown primary, anal, peritoneal, thyroid and unknown tumour streams. These are reflective of a patient population with limited treatment options. Tumour histology was most commonly adenocarcinoma (64.9%) followed by carcinoma (13.8%), squamous cell carcinoma (4.3%) and carcinoid (2.1%) (Table II). Other histology included papillary cancer, small cell carcinoma, serous and mucinous cystadenocarcinoma and glioblastoma. No patients were microsatellite instability high (MSI-H) (Table II). High tumour mutational burden (TMB-H) was found in 6.4% of patients (Table II).

**Table II:**
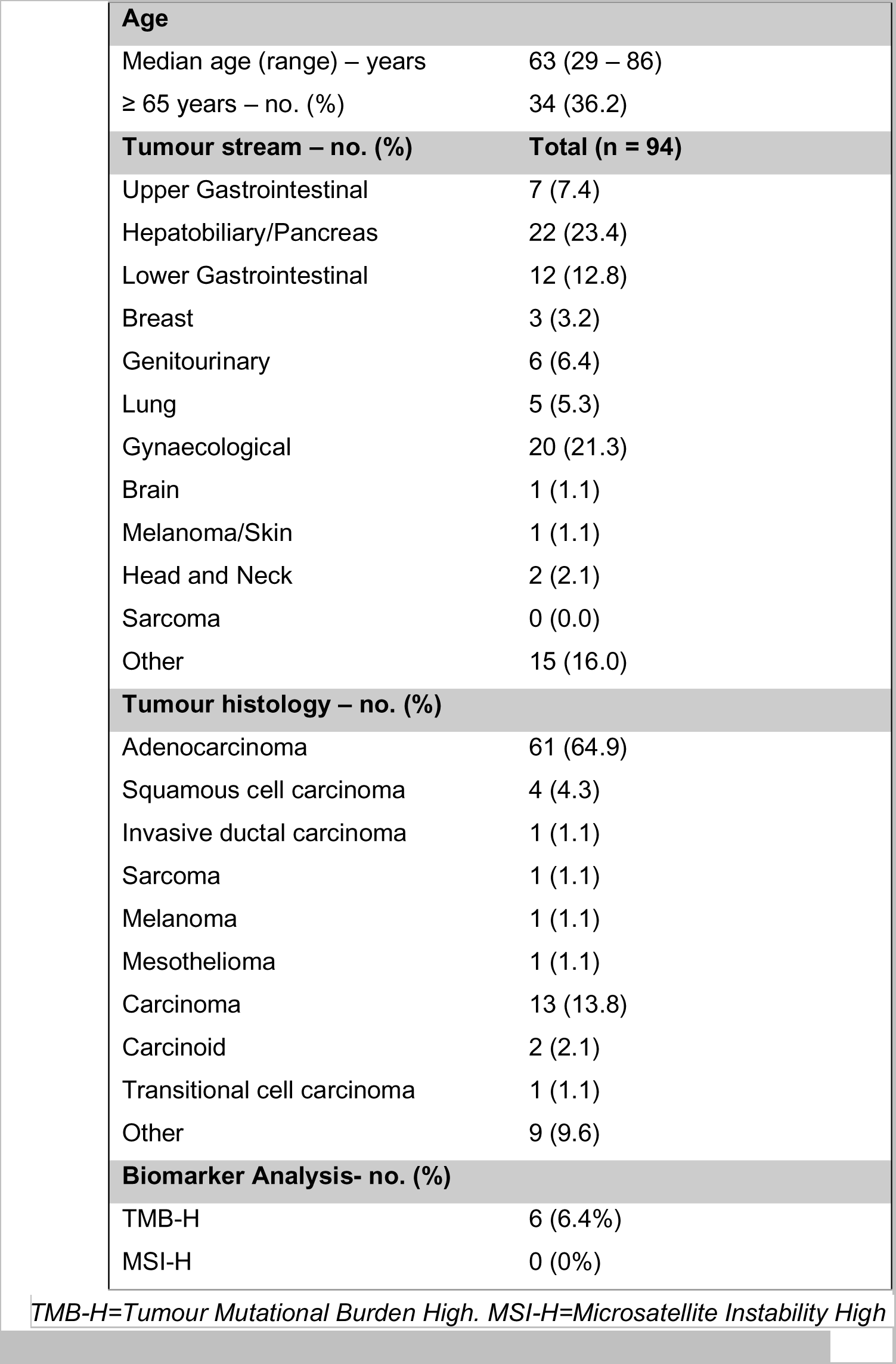
Demographics of patient cohort.

### Mutations

A total of 396 mutations were described from the 94 patient NGS reports, with 180 (45.4%) being gain of function mutations. The most common type of gain of function alteration was amplification (58.3%), with genes affected including *CCND1*, *FGF3*, *FGF19*, *FGF4*, *MYC*, *ERBB2* and *EGFR* affected most frequently (Figure 1). Non-synonymous single nucleotide variations (nsSNV) represented 35.6% of gain of function mutations (Figure 1). Genes most frequently altered were *KRAS*, *TP53*, *BRAF*, *PIK3CA* and *NRAS*. The frequency of fusions was 3.3% including *TMPRSS2*-*ERG*, *HNRNPH1*-*ETV4*, *ESR1*-*PLEKHG1* and *TPM3*-*ROS1*. Other alterations accounted for 2.7%.

**Figure 1:**
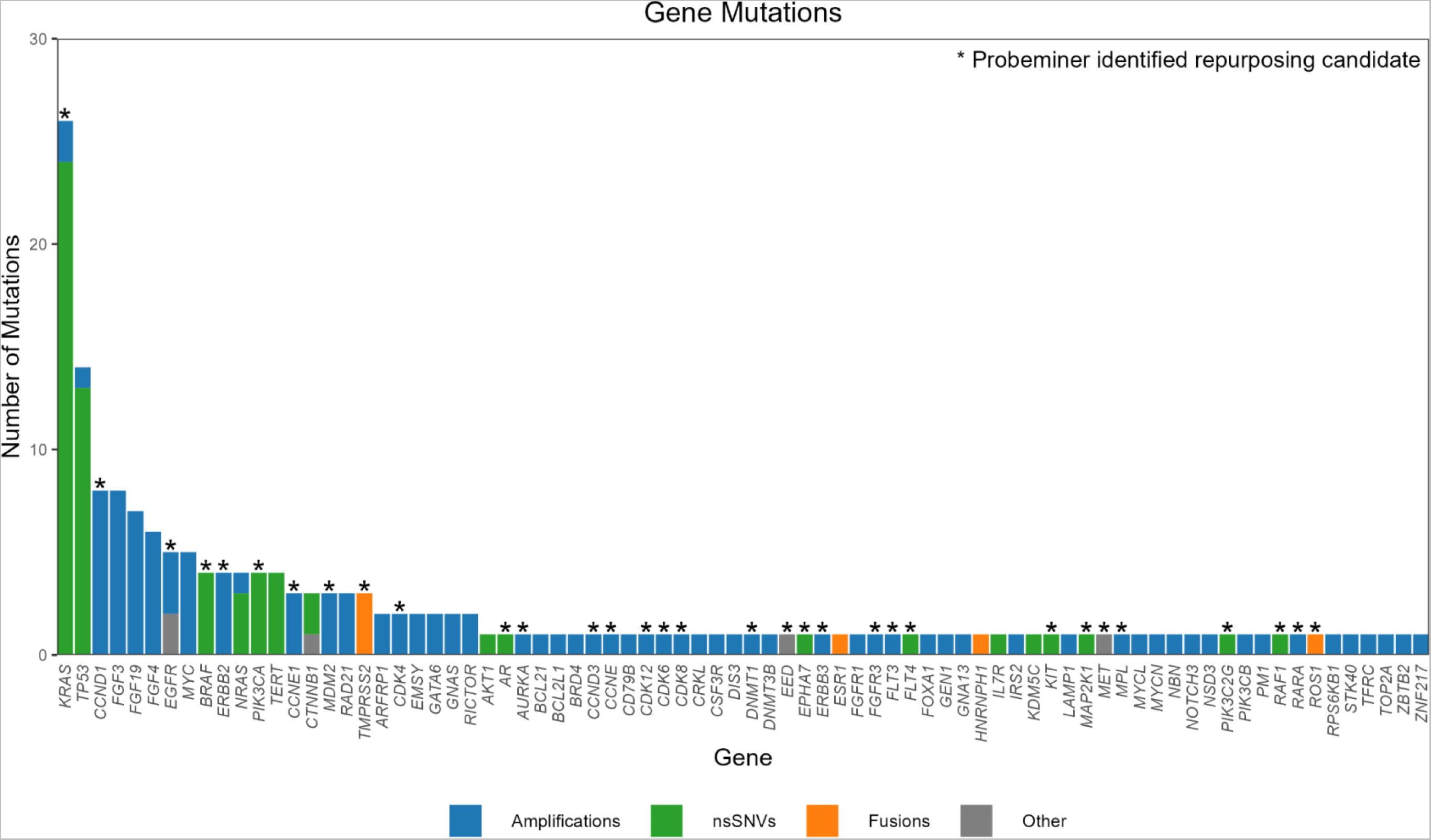
Histogram of gain of function mutations (n=180) Gain of function mutations included amplifications (blue), non-synonymous single-nucleotide variants (green), fusions (orange) and other mutations (grey). Asterisk indicates mutations for which a repurposing event was identified on Probeminer (n=32).

### Drug Repurposing

Seventy-five repurposing events were identified by Probe Miner, 80 by TOPOGRAPH and 50 by the Broad Institute DRH, which reduced to 32, 29 and 26 respectively after removing duplicate events (Figure 2). Within Probe Miner, there were 21 repurposing events with trial level evidence, 11 repurposing events without trial level evidence and four novel drug repurposing events (Figure 2). At a patient level, repurposing was applied to mutations for which no FDA-approved therapy or recruiting clinical trial was available (Figure 3). 13 unique patients had an off-label gene event (i.e., the repurposing approach identified a gene target for which there was no available clinical trial nationally).

**Figure 2:**
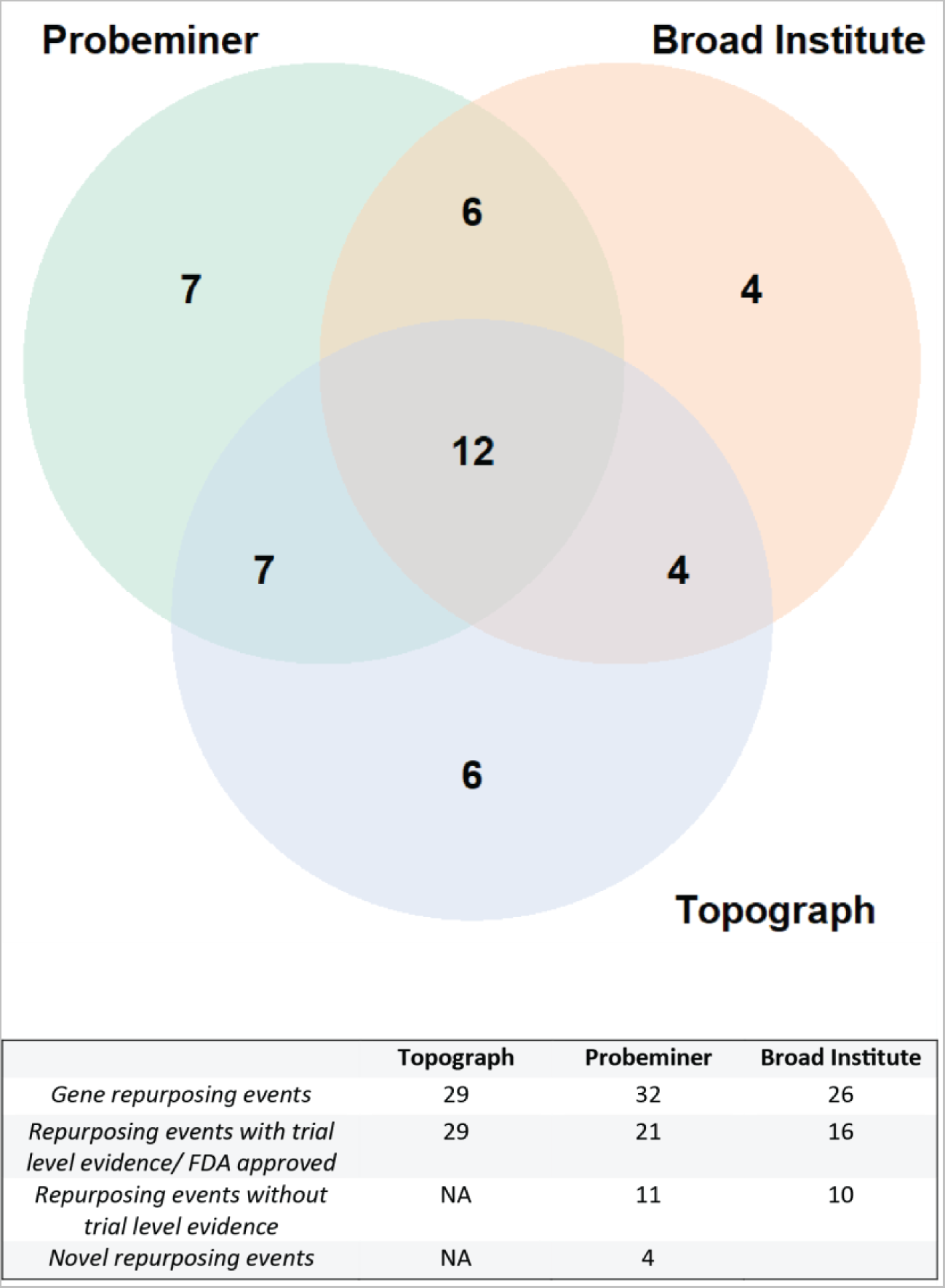
Repurposing events. Unique repurposing events identified via the three databases were evaluated for overlap as indicated in the Venn diagram (top). The breakdown of these repurposing events by trial level evidence and degree of novelty (bottom).

**Figure 3:**
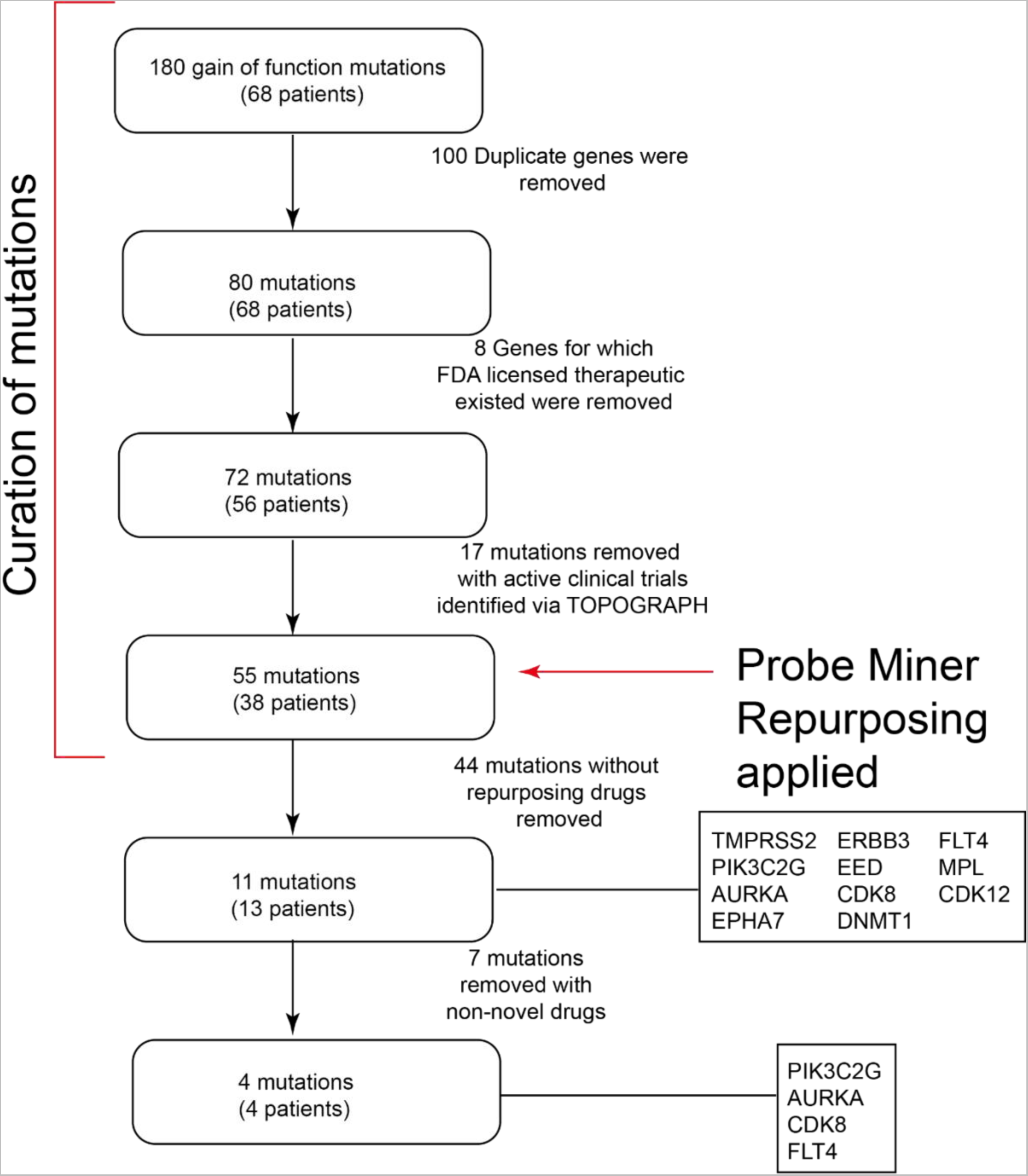
CONSORT diagram of selection of novel drug repurposing events, patient level. 180 gain of function mutations were identified across the cohort of patients, which represented 80 unique genes across 68 patients. Eight of these genes had FDA licensed therapies available and a further 17 had active, locally available clinical trials evaluating this biomarker. Of the 55 remaining mutations, 11 mutations (13 patients) had a repurposable drug identified by Probeminer or which 4 were considered novel.

The most common tumour type with targets found in Probe Miner was hepatobiliary/pancreatic (24% of patients), colorectal (18.7%), gynaecological (16%), genitourinary (9.3%) and upper gastrointestinal (5.3%). The most common types of mutations with probes found in Probe Miner were nsSNV (46.7%) and amplifications (46.7%) (Table III, Figure 1). The most common gene involved in nsSNV mutations was *KRAS* (25.3%), with *KRAS* G12D being most frequent. This was followed by *BRAF* (5.3%) and *PIK3CA* (5.3%) mutations. In terms of amplifications *CCND1* (10.7%), *CCNE1* (5.3%) and *ERBB2* (5.3%) were most involved. A full list of mutations identified is included in Supplementary Table III.

**Table III:** Mutations with drug probes found in Probeminer.

| TYPE OF MUTATION | MUTATED GENE | MUTATIONS (N) | MUTATIONS (%) |
| --- | --- | --- | --- |
| **NSSNV** | **TOTAL** | **35** | **46.7** |
|  | AR | 1 |  |
|  | BRAF | 4 | 5.3 |
|  | EED | 1 |  |
|  | EPHA7 | 1 |  |
|  | <u>FLT4*</u> | 1 |  |
|  | KIT | 1 |  |
|  | KRAS | 19 | 25.3 |
|  | MAP2K1 (MEK1) | 1 |  |
|  | <u>PIK3C2G*</u> | 1 |  |
|  | PIK3CA | 4 | 5.3 |
|  | RAF1 | 1 |  |
| **AMPLIFICATION** | **TOTAL** | **35** | **46.7** |
|  | <u>AURKA*</u> | 1 |  |
|  | CCND1 | 8 | 10.7 |
|  | CCND3 | 1 |  |
|  | CCNE1 | 4 | 5.3 |
|  | CDK12 | 1 |  |
|  | CDK4 | 2 |  |
|  | CDK6 | 1 |  |
|  | <u>CDK8*</u> | 1 |  |
|  | DNMT1 | 1 |  |
|  | EGFR | 5 |  |
|  | ERBB2 | 4 | 5.3 |
|  | ERBB3 | 1 |  |
|  | FGFR3 | 1 |  |
|  | FLT3 | 1 |  |
|  | <u>MDM2*</u> | 1 |  |
|  | MPL | 1 |  |
|  | RARA | 1 |  |
| **OTHER** | **TOTAL** | **1** | **1.3** |
|  | MET | 1 |  |
| **FUSION** | **TOTAL** | **4** | **5.3** |
|  | TMPRSS2-ERG | 3 |  |
|  | TPM3-ROS1 | 1 |  |
| **OVERALL TOTAL** |  | **75** |  |

\*Genes with novel drug repurposing probes found

### Novel Drug Repurposing Events

Within the Probe Miner drug events found, four novel repurposing targets were found (Figure 3), with multiple candidate drugs, of which the top two candidate repurposing events were further evaluated. Specific mutations included for *PIK3C2G* R1034H and *FLT4* V763M, whilst the other genes were amplified (Supplementary Table III). To demonstrate the validity of the drug repurposing events identified by Probe Miner, a comprehensive literature search was performed for the drug-target interactions considered novel. Strong biochemical data supported drug-target interactions in each case. For PIK3C2G-midostaurin, PIK3C2G-lapatinib, FLT4-axitinib, FLT4-tivozanib, AURKA-axitinib and CDK8-sorafenib experimental data demonstrating target selectivity was found (31, 32, 33). The potency was highest for FLT4-tivozanib (0.2 nM) and lowest for PIK3C2G-lapatinib (7,500 nM). Overall, literature supporting each drug-target interaction was demonstrated, with robust supporting biochemical data observable for all novel repurposing events.

Although Probe Miner is designed to evaluate drug-target binding, it is not designed to predict the functional consequences of this interaction. To evaluate whether prospective repurposing events are potentially functionally consequential, public datasets download from the Cancer Dependency Map Project were explored (24, 25, 26, 27, 34, 35, 36, 37). The Cancer Dependency Map Project builds upon the original Cancer Cell Line Encyclopedia (18), which involved the systematic molecular profiling of 1,000 cell lines, and performed large-scale functional genomics profiling via RNA-interference and CRISPR screens to identify genetic dependencies. In parallel, these cell lines had systematic drug sensitivity profiling performed. It was hypothesised that gene expression and gene-dependency in these public datasets would correlate with drug activity for each drug-target interaction in an exploratory analysis.

To evaluate the validity of this hypothesis, target gene expression (26) and CRISPR dependency (27) from the Cancer Dependency Map Project were plotted against drug sensitivity (log-fold change in cell viability), from either the Genomics of Drug Sensitivity Screens (38) or the Broad PRISM project (11) for all known FDA approved drug-target combinations (Supplementary Table I). An ANOVA was performed comparing the drug sensitivity with gene-expression or CRISPR dependency data analysed by quartiles. Of the FDA approved drug-target combinations, 55 had available drug sensitivity data, of which 23 (42%) had a statistically significant relationship between gene expression (n = 12) or gene dependency (n = 19) and drug sensitivity and 34 did not. Table IV lists the FDA drug-target combinations with statistically significant relationships between expression or CRISPR gene-dependency data and drug sensitivity and Figure 4 (top panels) demonstrate violin plots of the drug sensitivity data for osimertinib. Only eight drug-target combinations demonstrated a statistically significant relationship between both gene expression and CRISPR gene-dependency and drug sensitivity, all of which were drugs targeting EGFR or ERBB2, two of the best characterised oncogenes. Supplementary Table IV lists all FDA-approved drug-target combinations for which no statistically significant relationship was found between gene-expression/CRISPR gene-dependency data and drug sensitivity.

**Figure 4:**
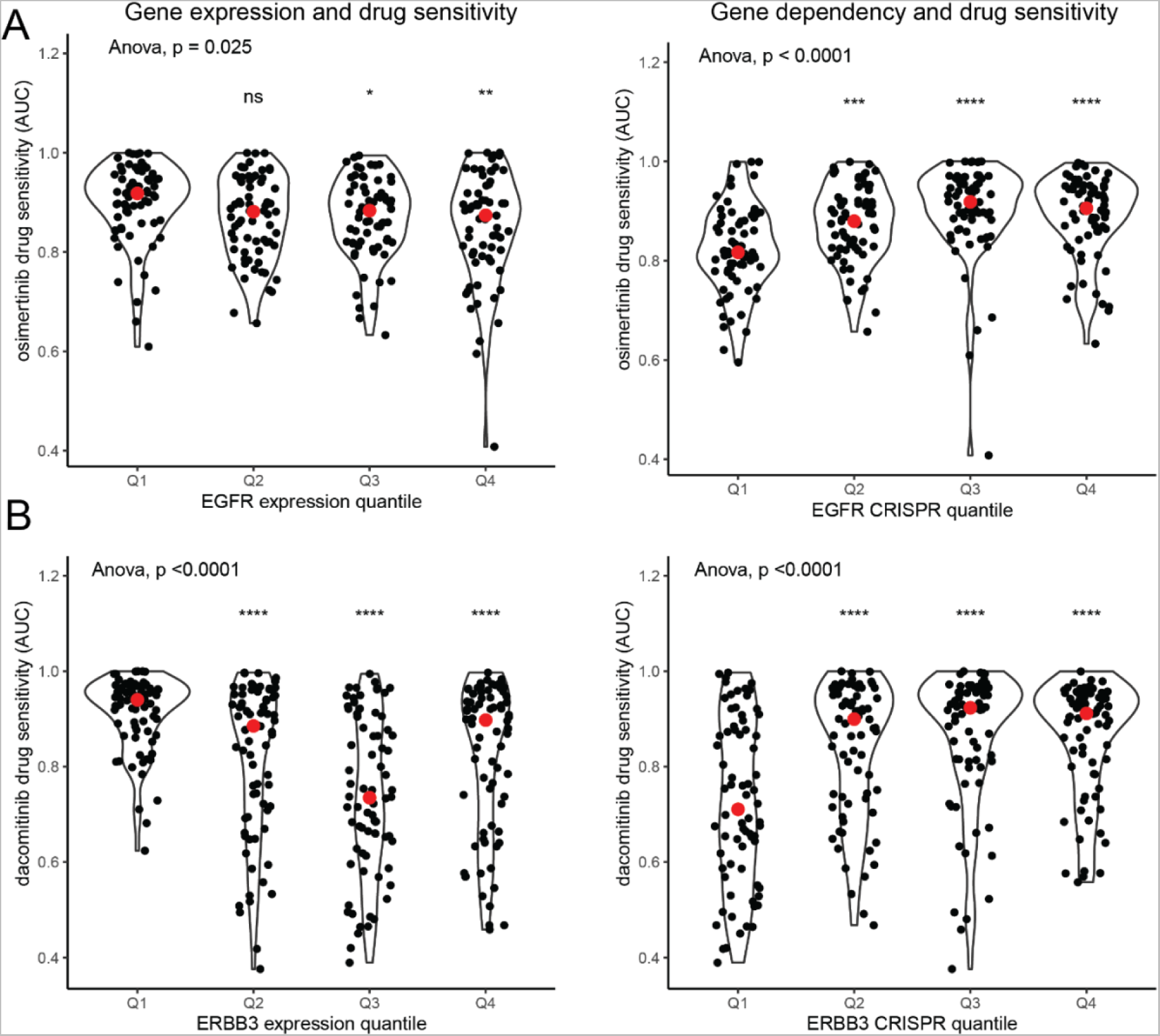
Gene expression and drug sensitivity. A) Drug sensitivity for osimertinib (area under the curve of a dose-response curve) according to EGFR expression (left) and EGFR gene-dependency (right). B) Drug sensitivity for dacomitinib (area under the curve of a dose-response curve) according to ERBB3 expression (left) and ERBB3 gene-dependency (right). Asterisks indicate results of t-tests comparing relevant quartile with the first quartile of gene expression/gene dependency: * p<0.05, **p<0.01, ***p<0.001, ****P<0.0001

**Table IV:**
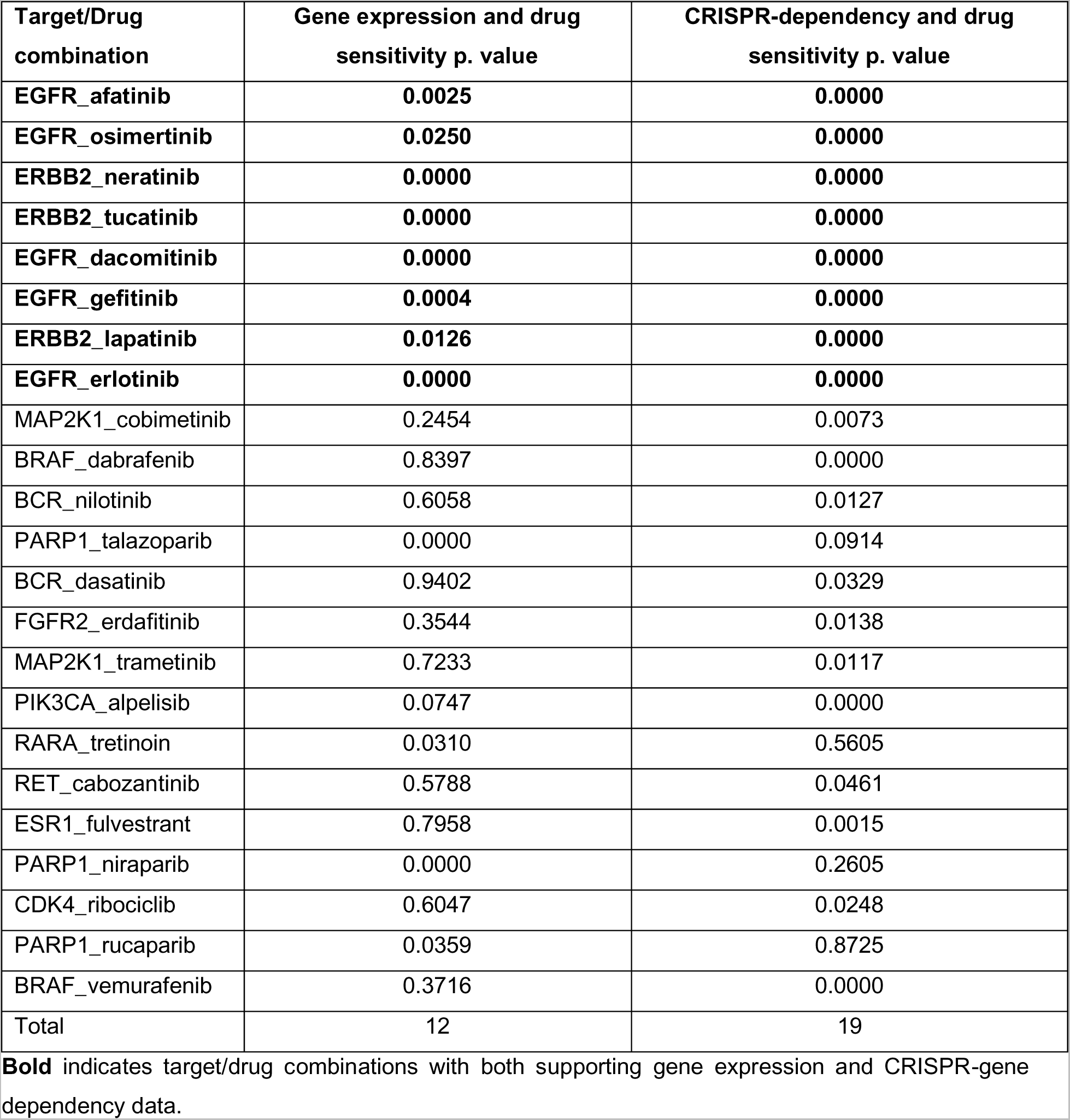
FDA approved drug-target combinations with supportive functional data from the Cancer Dependency Map Project.

**Table V:**
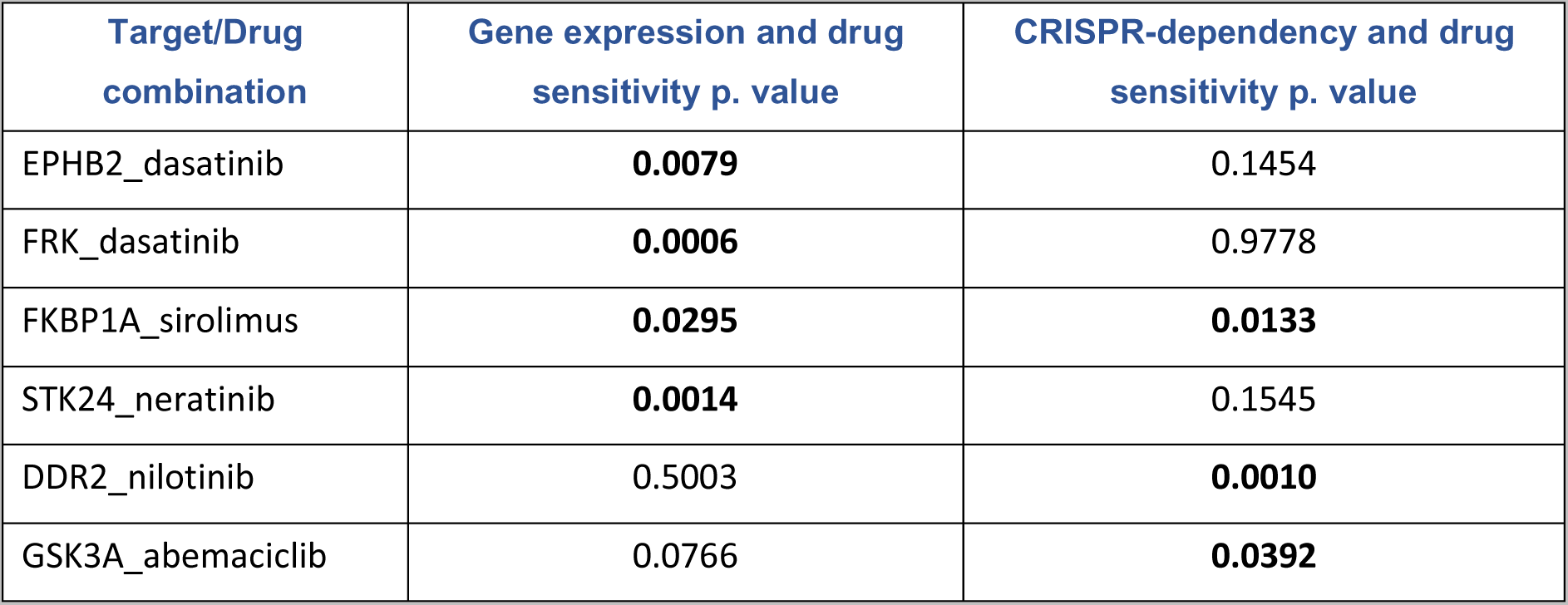
Probe Miner identified (global score > 0.7) repurposing drug-target combinations with supportive functional data from the Cancer Dependency Map Project.

To evaluate the functional consequences of potential repurposing events, target gene expression (26) and CRISPR dependency (27) from the Cancer Dependency Map Project were plotted against drug sensitivity (log-fold change in cell viability or area under the curve of a dose-response curve), from either the Genomics of Drug Sensitivity Screens (38) or the Broad PRISM project (11) for the 11 repurposing events identified by Probe Miner without trial level evidence. A statistically significant increased drug sensitivity was demonstrated with increased *PIK3C2G* expression and lapatinib (*p<0.001*),with *ERBB3* expression (*p<0.0001)* and *ERBB3* gene-dependency *(p<0.0001)* with multiple drugs including dacomitinib. Violin plots of the gene-expression and gene dependency data for dacomitinib-ERBB3 are shown in Figure 4 (bottom panels).

### Theoretical Drug Repurposing Events

The total repurposable genome was evaluated by analysing all gene targets with a Probe Miner global score >0.25 (from Probe Miner dataset) that were indexed as a licensed therapeutic in the Drugs@FDA database to get the total repurposable genome. All drug-gene combinations found in this analysis are listed in Supplementary Table IV A total of 1,968 theoretical repurposing events were identified. These genes were evaluated for frequency of mutations in the TCGA pan-cancer cohort (28) to assess the frequency of patients having possible mutations. 2,142 out of 2,922 (73%) samples had a mutation in a gene that was categorised as a theoretical repurposing event. Drug-gene combinations with a Probe Miner global score > 0.7 were further analysed as described above to identify potentially functionally consequential repurposing events. Six repurposing events with supportive functional data were found (Table V)(Supplementary Figure I).

## Discussion

A biomarker-driven precision oncology approach is increasingly used to enrich patients for clinical trials, however, trial access issues limit the utility of this approach for real-world patients. A drug repurposing approach, using established drugs with known safety profiles, potentially mitigates limitations of the current paradigm. Traditional drug repurposing relies upon serendipity and more recently systematic drug repurposing screens are being used for drug identification (11). Combining genomic biomarker testing with an *in silico* approach utilising pre-screened gene-drug interaction databases could enrich the identification of candidate drugs for repurposing to be formally tested in clinical trials.

In this study, using a real-world dataset of next generation sequencing molecular reports, we showed that a meaningful (14%) percentage of patients would have an additional off-label therapeutic identified by using a computational drug repurposing approach. This compares favourably with the results of the NCI-MATCH clinical trial, which found an actionable alteration rate of 38% (8). As our computational drug repurposing approach excludes mutations that would confer eligibility for local clinical trials, this additional incremental off-label therapeutic access is particularly meaningful. As only 17% of patients with actionable mutations identified on the NCI-MATCH clinical trial enrolled onto a sub-study (7), the computational drug repurposing approach may substantially expand the number of patients treatable with a biomarker-enriched approach. Overall, a repurposing rate of 14% was consistent with our hypothesis that a computational drug repurposing approach may identify novel therapeutic options for patients with no further access to standard therapies.

Several exploratory analyses were conducted which raised interesting findings that require further elucidation. Firstly, several drug-target interactions that have been previously elucidated in the medicinal chemistry literature, but are not well known, were identified. Preliminary exploratory functional analysis using publicly available datasets suggested that further validation of these targets may be warranted. Preclinical target validation is notoriously complex (39) and whilst these results are interesting, robust additional orthogonal validation is necessary to make further conclusions on the functional consequences of drug therapy.

As the pool of licensed therapeutics increases continually and only 11% of the kinome is currently characterised (15), and the targeted NGS panel used evaluated only 523 genes, successful validation of this approach could potentially meet a large unmet disease burden in oncology patients if the genome becomes better characterised and more extensive sequencing is performed. To explore the theoretical drug repurposing genome, all genes mapped to compounds with a Probe Miner global score >0.25 with an FDA-licensed therapeutic were evaluated for their frequency on the TCGA pan-cancer dataset, showing that 73% of samples in this cohort would have a theoretical drug repurposing event. This compares very favourably with the actionable alteration rate of 38% in the NCI-Match clinical trial.

There are several limitations to this study. In the main analysis, curation of variants of uncertain significance is fraught with difficulty. Extensive manual curation with a robust framework was performed to minimise the risk of mischaracterising mutations. Additionally, as Probe Miner is predominantly based on medicinal chemistry datasets that specifically assess drug-target binding, the assumption that drug-target binding results in meaningful anti-tumour activity is a large leap. Mitigating this, we evaluated Probe Miner’s ability to identify licensed FDA therapies linked to a biomarker, demonstrating high sensitivity (0.67-1.00) and specificity (0.99-1.00). These results strongly supported the validity of Probe Miner. Nevertheless, although strong inferences can be made about drug-target binding from this dataset, any conclusions regarding anti-tumour efficacy cannot be made from these data. Despite its inherent limitation, *in silico* functional predictions of utility of known therapies offer a novel strategy for rationally screening drug candidates to further examine in confirmatory phase 2 clinical trials.

In the exploratory analysis, robust mechanistic exploration of drug-target interactions was not performed. To make firm conclusions regarding possible anti-tumour activity, ideally *in vitro* cell viability assays with subsequent *in vivo* validation would be performed. For the theoretically repurposable drugs analysis, the cut-off Probe Miner global score of 0.25 was chosen as this was slightly greater than the lower bound of the range of global scores of FDA licensed therapies with known biomarkers. The Probe Miner global score is a relative score per gene target and using an absolute score of 0.25 as a cut-off is arbitrary. Additionally, many mutations annotated in the TCGA pan-cancer dataset are passenger alterations and do not contribute to oncogenesis; inclusion of such genomics findings could explain the higher-than-expected theoretical drug repurposing rate. Nevertheless, as the genome is further characterised and new therapies are licensed these numbers will only increase and a large baseline rate of theoretical drug repurposing events is fundamentally supportive of further research in this area.

## Conclusion

Despite significant advances in cancer care, patients with advanced solid tumours that are treatment refractory still face significant issues accessing novel therapeutics (8) and represent a large unmet need. Clinical drug development is extremely costly and drug repurposing can potentially substantially reduce drug development costs and timelines (9). We provide initial data demonstrating that in a real-world cohort of patients sequenced with a targeted NGS panel, that 14% of patients would have a possible, non-obvious drug repurposing candidate identified using a computational drug repurposing approach.

## Acknowledgements of research support

No financial support has been received.

There are no conflicts of interest which would influence the integrity of this manuscript.

This manuscript has been uploaded to BioRxiv prior to peer review and publication.

