## Supplementary Appendix for "Computational repurposing of oncology drugs through off-target drug binding interactions from pharmacological databases"

Supplementary Table I: Curated Table of Pharmacogenomics Biomarkers in Drug Labeling

| Drug | Biomarker | ENTREZ ID | Target | Tumour type | Chemical probe | Target combination license | Target tumour-specific | Target probe |
|---|---|---|---|---|---|---|---|---|
| **Abemaciclib** | ESR1 | 2099 | CDK4 | Breast | 1 | 0 | 1 | 0 |
| **Ado-trastuzumab emtansine** | ERBB2 | 2064 | ERBB2 | Breast | 0 | 0 | 0 | 1 |
| **Afatinib** | EGFR | 1956 | EGFR | Lung | 1 | 0 | 0 | 1 |
| **Alectinib** | ALK | 238 | ALK | Lung | 1 | 0 | 0 | 1 |
| **Alpelisib** | ERBB2 | 2064 | PIK3CA | Breast | 1 | 0 | 1 | 0 |
| **Amivantamab-vmjw** | EGFR | 1956 | EGFR | Lung | 0 | 0 | 0 | 1 |
| **Anastrozole** | ESR1 | 2099 | CYP19A1 | Breast | 1 | 0 | 1 | 1 |
| **Anastrozole** | PGR | 5241 | CYP19A1 | Breast | 1 | 0 | 1 | 1 |
| **Arsenic trioxide** | RARA | 5371 | RARA | PML | 1 | 0 | 1 | 0 |
| **Atezolizumab** | CD274 | 29126 | CD274 | Multiple | 0 | 0 | 0 | 1 |
| **Avapritinib** | KIT | 3815 | KIT | Multiple | 1 | 0 | 0 | 1 |
| **Avapritinib** | PDGFRA | 5156 | PDGFRA | Multiple | 1 | 0 | 0 | 1 |
| **Avelumab** | CD274 | 29126 | CD274 | Multiple | 0 | 0 | 0 | 1 |
| **Binimetinib** | BRAF | 673 | MAP2K1 | Melanoma | 1 | 1 | 0 | 0 |
| **Blinatumomab** | BCR | 613 | BCR | CML | 0 | 0 | 0 | 1 |
| **Blinatumomab** | CD19 | 930 | BCR | CML | 0 | 0 | 0 | 1 |
| **Bosutinib** | BCR | 613 | BCR | CML | 1 | 0 | 0 | 1 |
| **Brentuximab Vedotin** | ALK | 238 | TNFRSF8 | Lymphoma | 0 | 0 | 1 | 0 |
| **Brentuximab Vedotin** | TNFRSF8 | 943 | TNFRSF8 | Lymphoma | 0 | 0 | 0 | 1 |
| **Brigatinib** | ALK | 238 | ALK | Lung | 1 | 0 | 0 | 1 |
| **Busulfan** | BCR | 613 |  | Multiple | 1 | 0 | 1 | 1 |
| **Cabozantinib** | RET | 5979 | RET | Thyroid | 1 | 0 | 0 | 1 |
| **Capmatinib** | MET | 4233 | MET | Lung | 1 | 0 | 0 | 1 |
| **Cemiplimab-rwlc** | CD274 | 29126 | CD274 | Multiple | 0 | 0 | 0 | 1 |
| **Ceritinib** | ALK | 238 | ALK | Lung | 1 | 0 | 0 | 1 |
| **Cetuximab** | EGFR | 1956 | EGFR | Colorectal | 0 | 0 | 0 | 1 |
| **Cobimetinib** | BRAF | 673 | MAP2K1 | Melanoma | 1 | 1 | 0 | 0 |
| **Crizotinib** | ALK | 238 | ALK | Lung | 1 | 0 | 0 | 1 |
| **Crizotinib** | ROS1 | 6098 | ROS1 | Lung | 1 | 0 | 0 | 1 |
| **Dabrafenib** | BRAF | 673 | BRAF | Multiple | 1 | 0 | 0 | 1 |
| **Dabrafenib** | KRAS | 3845 | BRAF | Multiple | 1 | 0 | 1 | 0 |
| **Dacomitinib** | EGFR | 1956 | EGFR | Lung | 1 | 0 | 0 | 1 |
| **Dasatinib** | BCR | 1956 | BCR | CML | 1 | 0 | 0 | 1 |
| **Denileukin Diftitox** | IL2RA | 3559 | IL2RA | CTCL | 0 | 0 | 0 | 1 |
| **Dinutuximab** | MYCN | 4613 | GD2 | Neuroblastoma | 0 | 0 | 1 | 1 |
| **Dostarlimab-gxly** | MLH1 | 4292 | CD274 | Multiple | 0 | 0 | 0 | 0 |
| **Dostarlimab-gxly** | MSH2 | 4436 | CD274 | Multiple | 0 | 0 | 0 | 0 |
| **Dostarlimab-gxly** | MSH6 | 2956 | CD274 | Multiple | 0 | 0 | 0 | 0 |

| | | | | | | | | |
|---|---|---|---|---|---|---|---|---|
| **Dostarlimab-gxly** | PMS2 | 5395 | CD274 | Multiple | 0 | 0 | 0 | 0 |
| **Durvalumab** | CD274 | 29126 | CD274 | Multiple | 0 | 0 | 0 | 1 |
| **Enasidenib** | IDH2 | 3418 | IDH2 | Multiple | 1 | 0 | 0 | 1 |
| **Encorafenib** | BRAF | 673 | BRAF | Multiple | 1 | 0 | 0 | 1 |
| **Encorafenib** | KRAS | 3845 | BRAF | Multiple | 1 | 1 | 0 | 0 |
| **Enfortumab Vedotin-ejfv** | NECTIN4 | 81607 | Nectin4 | Urothelial | 0 | 0 | 0 | 1 |
| **Entrectinib** | NTRK1 | 4914 | NTRK1 | Multiple | 1 | 0 | 0 | 1 |
| **Entrectinib** | NTRK2 | 4915 | NTRK2 | Multiple | 1 | 0 | 0 | 1 |
| **Entrectinib** | NTRK3 | 4916 | NTRK3 | Multiple | 1 | 0 | 0 | 1 |
| **Entrectinib** | ROS1 | 6098 | ROS1 | Lung | 1 | 0 | 0 | 1 |
| **Erdafitinib** | FGFR2 | 2263 | FGFR2 | Biliary | 1 | 0 | 0 | 1 |
| **Erdafitinib** | FGFR3 | 2261 | FGFR3 | Biliary | 1 | 0 | 0 | 1 |
| **Erlotinib** | EGFR | 1956 | EGFR | Lung | 1 | 0 | 0 | 1 |
| **Everolimus** | ERBB2 | 2064 | MTOR | Breast | 1 | 1 | 1 | 0 |
| **Exemestane** | ESR1 | 2099 | CYP19A1 | Breast | 1 | 0 | 1 | 1 |
| **Exemestane** | PGR | 5241 | CYP19A1 | Breast | 1 | 0 | 1 | 1 |
| **Fam-Trastuzumab Deruxtecan-nxki** | ERBB2 | 2064 | ERBB2 | Breast | 0 | 0 | 0 | 1 |
| **Fulvestrant** | ESR1 | 2099 | ESR1 | Breast | 1 | 0 | 1 | 1 |
| **Fulvestrant** | PGR | 5241 | ESR1 | Breast | 1 | 0 | 1 | 1 |
| **Gefitinib** | EGFR | 1956 | EGFR | Lung | 1 | 0 | 0 | 1 |
| **Gemtuzumab Ozogamicin** | CD33 | 945 | CD33 | AML | 0 | 0 | 0 | 1 |
| **Gilteritinib** | FLT3 | 2322 | FLT3 | AML | 1 | 0 | 0 | 1 |
| **Goserelin** | ESR1 | 2099 | GNRHR | Breast | 1 | 0 | 1 | 0 |
| **Goserelin** | PGR | 5241 | GNRHR | Breast | 1 | 0 | 1 | 0 |
| **Ibrutinib** | MYD88 | 4615 | BTK | Multiple | 1 | 0 | 0 | 0 |
| **Imatinib** | BCR | 613 | BCR | CML | 1 | 0 | 0 | 1 |
| **Imatinib** | PDGFRA | 5156 | | | 1 | 0 | 0 | 1 |
| **Imatinib** | KIT | 3815 | KIT | Multiple | 1 | 0 | 0 | 1 |
| **Imatinib** | PDGFRB | 5159 | | | 1 | 0 | 0 | 1 |
| **Infigratinib** | FGFR2 | 2263 | FGFR2 | Biliary | 1 | 0 | 0 | 1 |
| **Inotuzumab Ozogamicin** | BCR | 613 | CD22 | ALL | 0 | 0 | 0 | 1 |
| **Ipilimumab** | MLH1 | 4292 | CTLA4 | Multiple | 0 | 1 | 0 | 0 |
| **Ipilimumab** | MSH2 | 4436 | CTLA4 | Multiple | 0 | 1 | 0 | 0 |
| **Ipilimumab** | MSH6 | 2956 | CTLA4 | Multiple | 0 | 1 | 0 | 0 |
| **Ipilimumab** | PMS2 | 5395 | CTLA4 | Multiple | 0 | 1 | 0 | 0 |
| **Ivosidenib** | IDH1 | 3417 | IDH1 | Multiple | 1 | 0 | 0 | 1 |
| **Ixabepilone** | ERBB2 | 2064 | | Breast | 1 | 1 | 1 | 0 |
| **Ixabepilone** | ESR1 | 2099 | | Breast | 1 | 0 | 1 | 0 |
| **Ixabepilone** | PGR | 5241 | | Breast | 1 | 0 | 1 | 0 |
| **Lapatinib** | ERBB2 | 2064 | ERBB2 | Breast | 1 | 0 | 0 | 1 |
| **Lapatinib** | ESR1 | 2099 | ERBB2 | Breast | 1 | 0 | 1 | 0 |
| **Lapatinib** | PGR | 5241 | ERBB2 | Breast | 1 | 0 | 1 | 0 |
| **Larotrectinib** | NTRK1 | 4914 | NTRK1 | Multiple | 1 | 0 | 0 | 1 |

| | | | | | | | | |
|---|---|---|---|---|---|---|---|---|
| **Larotrectinib** | NTRK2 | 4915 | NTRK2 | Multiple | 1 | 0 | 0 | 1 |
| **Larotrectinib** | NTRK3 | 4916 | NTRK3 | Multiple | 1 | 0 | 0 | 1 |
| **Lenvatinib** | MLH1 | 4292 | KDR | Multiple | 1 | 1 | 0 | 0 |
| **Lenvatinib** | MSH2 | 4436 | KDR | Multiple | 1 | 1 | 0 | 0 |
| **Lenvatinib** | MSH6 | 2956 | KDR | Multiple | 1 | 1 | 0 | 0 |
| **Lenvatinib** | PMS2 | 5395 | KDR | Multiple | 1 | 1 | 0 | 0 |
| **Letrozole** | ESR1 | 2099 | CYP19A1 | Breast | 1 | 0 | 1 | 0 |
| **Letrozole** | PGR | 5241 | CYP19A1 | Breast | 1 | 0 | 1 | 0 |
| **Lorlatinib** | ROS1 | 6098 | ROS1 | Lung | 1 | 0 | 0 | 1 |
| **Lutetium Dotatae Lu-177** | SSTR1 | 6751 | | | 0 | 0 | 0 | 1 |
| **Margetuximab-cmkb** | ERBB2 | 2064 | ERBB2 | Breast | 0 | 0 | 0 | 1 |
| **Margetuximab-cmkb** | FCGR2A | 2212 | ERBB2 | Breast | 0 | 1 | 0 | 1 |
| **Margetuximab-cmkb** | FCGR2B | 2213 | ERBB2 | Breast | 0 | 1 | 0 | 1 |
| **Margetuximab-cmkb** | FCGR3A | 2214 | ERBB2 | Breast | 0 | 1 | 0 | 1 |
| **Midostaurin** | FLT3 | 2322 | FLT3 | AML | 1 | 0 | 0 | 1 |
| **Midostaurin** | KIT | 3815 | FLT3 | AML | 1 | 0 | 0 | 1 |
| **Midostaurin** | NPM1 | 4869 | FLT3 | AML | 1 | 0 | 0 | 1 |
| **Neratinib** | ESR1 | 2099 | ERBB2 | Breast | 1 | 0 | 1 | 0 |
| **Neratinib** | PGR | 5241 | ERBB2 | Breast | 1 | 0 | 1 | 0 |
| **Nilotinib** | BCR | 613 | BCR | CML | 1 | 0 | 0 | 1 |
| **Niraparib** | BRCA1 | 672 | PARP1 | Multiple | 1 | 0 | 0 | 1 |
| **Niraparib** | BRCA2 | 675 | PARP1 | Multiple | 1 | 0 | 0 | 1 |
| **Nivolumab** | CD274 | 29126 | CD274 | Multiple | 0 | 0 | 0 | 1 |
| **Nivolumab** | MLH1 | 4292 | CD274 | Multiple | 0 | 0 | 0 | 0 |
| **Nivolumab** | MSH2 | 4436 | CD274 | Multiple | 0 | 0 | 0 | 0 |
| **Nivolumab** | MSH6 | 2956 | CD274 | Multiple | 0 | 0 | 0 | 0 |
| **Nivolumab** | PMS2 | 5395 | CD274 | Multiple | 0 | 0 | 0 | 0 |
| **Olaparib** | BRCA1 | 672 | PARP1 | Multiple | 1 | 0 | 0 | 0 |
| **Olaparib** | BRCA2 | 675 | PARP1 | Multiple | 1 | 0 | 0 | 0 |
| **Olaparib** | ESR1 | 2099 | PARP1 | Multiple | 1 | 0 | 1 | 0 |
| **Olaparib** | PGR | 5241 | PARP1 | Multiple | 1 | 0 | 1 | 0 |
| **Olaparib** | PPP2R2A | 5520 | PARP1 | Multiple | 1 | 0 | 0 | 0 |
| **Olaratumab** | PDGFRA | 5156 | PDGFRA | Sarcoma | 0 | 0 | 0 | 1 |
| **Omacetaxine** | BCR | 613 | | CML | 1 | 0 | 0 | 1 |
| **Osimertinib** | EGFR | 1956 | EGFR | Lung | 1 | 0 | 0 | 1 |
| **Palbociclib** | ESR1 | 2099 | CDK4 | Breast | 1 | 0 | 1 | 0 |
| **Panitumumab** | EGFR | 1956 | EGFR | Colorectal | 0 | 0 | 0 | 1 |
| **Pembrolizumab** | CD274 | 29126 | CD274 | Multiple | 0 | 0 | 0 | 1 |
| **Pembrolizumab** | MLH1 | 4292 | CD274 | Multiple | 0 | 0 | 0 | 0 |
| **Pembrolizumab** | MSH2 | 4436 | CD274 | Multiple | 0 | 0 | 0 | 0 |
| **Pembrolizumab** | MSH6 | 2956 | CD274 | Multiple | 0 | 0 | 0 | 0 |
| **Pembrolizumab** | PMS2 | 5395 | CD274 | Multiple | 0 | 0 | 0 | 0 |
| **Pemigatinib** | FGFR2 | 2263 | FGFR2 | Biliary | 1 | 0 | 0 | 1 |
| **Pertuzumab** | ERBB2 | 2064 | ERBB2 | Breast | 0 | 0 | 0 | 1 |

| | | | | | | | | |
|---|---|---|---|---|---|---|---|---|
| **Ponatinib** | BCR | 613 | BCR | CML | 1 | 0 | 0 | 1 |
| **Pralsetinib** | RET | 5979 | RET | Lung | 1 | 0 | 0 | 1 |
| **Raloxifene** | ESR1 | 2099 | ESR1 | Breast | 1 | 0 | 0 | 1 |
| **Ramucirumab** | EGFR | 1956 | EGFR | Multiple | 0 | 0 | 0 | 0 |
| **Regorafenib** | KRAS | 3845 | KDR | Colorectal | 1 | 0 | 1 | 0 |
| **Ribociclib** | ESR1 | 2099 | CDK4 | Breast | 1 | 0 | 1 | 0 |
| **Ribociclib** | PGR | 5241 | CDK4 | Breast | 1 | 0 | 1 | 0 |
| **Rituximab** | MS4A1 | 931 | CD20 | Multiple | 0 | 0 | 1 | 0 |
| **Rucaparib** | BRCA1 | 672 | PARP1 | Multiple | 1 | 0 | 0 | 0 |
| **Rucaparib** | BRCA2 | 675 | PARP1 | Multiple | 1 | 0 | 0 | 0 |
| **Sotorasib** | KRAS | 3845 | KRAS | Multiple | 1 | 0 | 0 | 1 |
| **Talazoparib** | ERBB2 | 2064 | PARP1 | Multiple | 1 | 0 | 1 | 0 |
| **Tamoxifen** | ESR1 | 2099 | ESR1 | Breast | 1 | 0 | 1 | 1 |
| **Tamoxifen** | PGR | 5241 | ESR1 | Breast | 1 | 0 | 1 | 0 |
| **Tepotinib** | ALK | 238 | MET | Lung | 1 | 0 | 0 | 0 |
| **Tepotinib** | EGFR | 1956 | MET | Lung | 1 | 0 | 0 | 0 |
| **Tepotinib** | MET | 4233 | MET | Lung | 1 | 0 | 0 | 1 |
| **Toremifene** | ESR1 | 2099 | ESR1 | Breast | 1 | 0 | 1 | 1 |
| **Trametinib** | BRAF | 673 | MAP2K1 | Melanoma | 1 | 1 | 0 | 0 |
| **Trastuzumab** | ERBB2 | 2064 | ERBB2 | Breast | 0 | 0 | 0 | 1 |
| **Tretinoin** | RARA | 5371 | RARA | PML | 1 | 0 | 0 | 1 |
| **Tucatinib** | ERBB2 | 2064 | ERBB2 | Breast | 1 | 0 | 0 | 1 |
| **Vemurafenib** | BRAF | 673 | BRAF | Multiple | 1 | 0 | 0 | 1 |
| **Venetoclax** | FLT3 | 2322 | BCL2 | Multiple | 1 | 0 | 1 | 0 |
| **Venetoclax** | IDH1 | 3417 | BCL2 | Multiple | 1 | 0 | 1 | 0 |
| **Venetoclax** | IDH2 | 3418 | BCL2 | Multiple | 1 | 0 | 1 | 0 |
| **Venetoclax** | IGH | 3492 | BCL2 | Multiple | 1 | 0 | 1 | 0 |
| **Venetoclax** | NPM1 | 4869 | BCL2 | Multiple | 1 | 0 | 1 | 0 |
| **Venetoclax** | TP53 | 7157 | BCL2 | Multiple | 1 | 0 | 1 | 0 |

Supplementary Table II: Probeminer accuracy for FDA approved drugs

| **Gene** | **Drug** | **Probe Miner score** | **Rank** |
|---|---|---|---|
| BCR | ponatinib | 0.8 | 1 |
| NTRK1 | entrectinib | 0.79 | 1 |
| PIK3CA | alpelisib | 0.78 | 1 |
| CDK4 | palbociclib | 0.76 | 1 |
| MTOR | everolimus | 0.74 | 1 |
| CDK4 | abemaciclib | 0.73 | 2 |
| MET | capmatinib | 0.72 | 2 |
| EGFR | afatinib | 0.71 | 3 |
| MAP2K1 | cobimetinib | 0.67 | 3 |
| NTRK3 | entrectinib | 0.66 | 1 |
| ESR1 | raloxifene | 0.65 | 6 |
| NTRK2 | entrectinib | 0.65 | 1 |

| | | | |
|---|---|---|---|
| BCR | imatinib | 0.64 | 1 |
| ROS1 | crizotinib | 0.64 | 2 |
| EGFR | dacomitinib | 0.63 | 10 |
| MET | tepotinib | 0.63 | 10 |
| BTK | ibrutinib | 0.63 | 3 |
| EGFR | osimertinib | 0.61 | 33 |
| BRAF | vemurafenib | 0.61 | 3 |
| BRAF | encorafenib | 0.6 | 4 |
| KDR | lenvatinib | 0.6 | 26 |
| KDR | regorafenib | 0.6 | 27 |
| MAP2K1 | trametinib | 0.6 | 10 |
| ALK | ceritinib | 0.6 | 11 |
| RET | cabozantinib | 0.59 | 10 |
| RARA | tretinoin | 0.59 | 8 |
| ERBB2 | neratinib | 0.58 | 29 |
| PARP1 | olaparib | 0.58 | 3 |
| BRAF | dabrafenib | 0.58 | 7 |
| ROS1 | entrectinib | 0.57 | 7 |
| CYP19A1 | letrozole | 0.57 | 5 |
| ESR1 | fulvestrant | 0.56 | 61 |
| PARP1 | niraparib | 0.56 | 4 |
| FGFR2 | infigratinib | 0.55 | 8 |
| PARP1 | talazoparib | 0.54 | 5 |
| MAP2K1 | binimetinib | 0.54 | 19 |
| ALK | brigatinib | 0.54 | 26 |
| BCR | bosutinib | 0.53 | 35 |
| NTRK1 | larotrectinib | 0.51 | 20 |
| ESR1 | tamoxifen | 0.48 | 171 |
| FLT3 | gilteritinib | 0.48 | 161 |
| ERBB2 | lapatinib | 0.44 | 124 |
| CYP19A1 | anastrozole | 0.44 | 48 |
| CYP19A1 | exemestane | 0.4 | 77 |
| BCR | imatinib | 0.39 | 101 |
| EGFR | erlotinib | 0.38 | 1135 |
| BCL2 | venetoclax | 0.38 | 116 |
| IDH2 | enasidenib | 0.38 | 10 |
| BCR | nilotinib | 0.37 | 127 |
| BCR | dasatinib | 0.37 | 129 |
| PDGFRA | avapritinib | 0.36 | 233 |
| EGFR | gefitinib | 0.33 | 1977 |
| ROS1 | lorlatinib | 0.33 | 59 |
| KIT | avapritinib | 0.33 | 410 |
| KIT | imatinib | 0.32 | 472 |
| FGFR2 | erdafitinib | 0.32 | 575 |
| FGFR3 | erdafitinib | 0.32 | 349 |
| ALK | crizotinib | 0.31 | 482 |

| | | | |
|---|---|---|---|
| FLT3 | midostaurin | 0.3 | 1030 |
| PARP1 | rucaparib | 0.29 | 458 |
| NTRK3 | larotrectinib | 0.25 | 89 |
| NTRK2 | larotrectinib | 0.25 | 90 |
| IDH1 | ivosidenib | 0.19 | 491 |
| ALK | alectinib | NA | NA |
| CDK4 | ribociclib | NA | NA |
| ESR1 | toremifene | NA | NA |
| ERBB2 | tucatinib | NA | NA |

Supplementary Table III: Types of gain of function non-synonymous single nucleotide variations and amplification mutations

| TYPE OF GAIN OF FUNCTION MUTATION | MUTATED GENE | MUTATION | MUTATIONS (No.) | % OF MUTATIONS |
|---|---|---|---|---|
| **AMPLIFICATION** | | **TOTAL** | **105** | **58.3** |
| | ARFRP1 | | 2 | |
| | AURKA | | 1 | |
| | BCL21 | | 1 | |
| | BCL2L1 | | 1 | |
| | BRD4 | | 1 | |
| | C11orf30 (EMSY) | | 2 | |
| | CCND1 | | 8 | |
| | CCND3 | | 1 | |
| | CCNE1 | | 4 | |
| | CD79B | | 1 | |
| | CDK12 | | 1 | |
| | CDK4 | | 2 | |
| | CDK6 | | 1 | |
| | CDK8 | | 1 | |
| | CRKL | | 1 | |
| | CSF3R | | 1 | |
| | DIS3 | | 1 | |
| | DNMT1 | | 1 | |
| | DNMT3B | | 1 | |
| | EGFR | | 3 | |
| | ERBB2 | | 4 | |
| | ERBB3 | | 1 | |
| | FGF19 | | 7 | |
| | FGF3 | | 8 | |
| | FGF4 | | 6 | |
| | FGFR1 | | 1 | |
| | FGFR3 | | 1 | |
| | FLT3 | | 1 | |
| | FOXA1 | | 1 | |
| | GATA6 | | 2 | |
| | GEN1 | | 1 | |
| | GNA13 | | 1 | |
| | GNAS | | 2 | |
| | IRS2 | | 1 | |
| | KRAS | | 2 | |
| | LAMP1 | | 1 | |
| | MDM2 | | 3 | |
| | MPL | | 1 | |
| | MYC | | 5 | |
| | MYCL | | 1 | |

| | | | | | |
|---|---|---|---|---|---|
| | MYCN | | | 1 | |
| | NBN | | | 1 | |
| | NOTCH3 | | | 1 | |
| | NRAS | | | 1 | |
| | NSD3 | | | 1 | |
| | PIK3CB | | | 1 | |
| | PM1 | | | 1 | |
| | RAD21 | | | 3 | |
| | RARA | | | 1 | |
| | RICTOR | | | 2 | |
| | RPS6KB1 | | | 1 | |
| | STK40 | | | 1 | |
| | TFRC | | | 1 | |
| | TOP2A | | | 1 | |
| | TP53 | | | 1 | |
| | ZBTB2 | | | 1 | |
| | ZNF217 | | | 1 | |
| **nsSNV** | | **TOTAL** | | **64** | **35.6** |
| | AKT1 | E17K | | 1 | |
| | AR | L702H | | 1 | |
| | BRAF | | | 4 | |
| | | L587R | 1 | | |
| | | N581I | 1 | | |
| | | N581T | 1 | | |
| | | V600E | 1 | | |
| | CTNNB1 | | | 2 | |
| | EPHA7 | c.97+2T>C | | 1 | |
| | FLT4 | V763M | | 1 | |
| | IL7R | K395R | | 1 | |
| | KDM5C | S717L | | 1 | |
| | KIT | D816E | | 1 | |
| | KRAS | | | 24 | |
| | | G12D | 13 | | |
| | | Q61H | 1 | | |
| | | A146T | 1 | | |
| | | G12C | 2 | | |
| | | G12S | 1 | | |
| | | G12V | 4 | | |
| | | G13D | 1 | | |
| | | G12R | 1 | | |
| | MAP2K1 (MEK1) | K57N | | 1 | |
| | NRAS | | | 3 | |
| | | G12C | 1 | | |
| | | Q61R | 2 | | |
| | PIK3C2G | R1034H | | 1 | |
| | PIK3CA | | | 4 | |
| | | E542K | 2 | | |
| | | H1047R | 2 | | |

| | | | |
|---|---|---|---|
| | RAF1 | S257L | 1 |
| | TERT promoter | | 4 |
| | TP53 | | 13 |

Supplementary Table IV: Total repurposable genome

| Drug | Gene | Probeminer score |
|---|---|---|
| **axitinib** | FLT1 | 0.86 |
| axitinib | FLT4 | 0.86 |
| axitinib | KDR | 0.84 |
| tretinoin | RXRG | 0.84 |
| dasatinib | EPHA5 | 0.84 |
| dasatinib | EPHB4 | 0.83 |
| dacomitinib | ERBB3 | 0.82 |
| dasatinib | ABL2 | 0.82 |
| dasatinib | EPHA4 | 0.80 |
| dasatinib | EPHA3 | 0.80 |
| bosutinib | MAP4K5 | 0.80 |
| diethylstilbestrol | ESR2 | 0.80 |
| ponatinib | ABL1 | 0.80 |
| axitinib | AURKC | 0.80 |
| bosutinib | LCK | 0.80 |
| dasatinib | EPHB2 | 0.79 |
| idelalisib | PIK3CD | 0.79 |
| zafirlukast | CYSLTR1 | 0.79 |
| dasatinib | EPHA8 | 0.78 |
| afatinib | ERBB4 | 0.78 |
| dasatinib | EPHA2 | 0.78 |
| dasatinib | FRK | 0.78 |
| alpelisib | PIK3CA | 0.78 |
| indomethacin | AR | 0.78 |
| tofacitinib | JAK3 | 0.77 |
| dasatinib | EPHB1 | 0.77 |
| dasatinib | LYN | 0.77 |
| dasatinib | FGR | 0.77 |
| gefitinib | GAK | 0.76 |
| finasteride | SRD5A2 | 0.76 |
| regorafenib | EPHX2 | 0.76 |
| neratinib | STK25 | 0.76 |
| dasatinib | FYN | 0.76 |
| ibrutinib | BLK | 0.76 |
| sirolimus | FKBP1A | 0.76 |
| palbociclib | CDK4 | 0.76 |
| axitinib | PLK4 | 0.76 |
| ribociclib | CCND3 | 0.76 |
| fedratinib | DAPK3 | 0.75 |
| neratinib | STK24 | 0.75 |
| palbociclib | CCND2 | 0.75 |
| palbociclib | CDK6 | 0.75 |
| ibrutinib | BMX | 0.75 |

| | | |
|---|---|---|
| erlotinib | MAP3K19 | 0.75 |
| sorafenib | DDR1 | 0.75 |
| nilotinib | DDR2 | 0.75 |
| fluphenazine | DRD2 | 0.74 |
| sunitinib | MYLK4 | 0.74 |
| ibrutinib | HCK | 0.74 |
| dasatinib | SRC | 0.74 |
| diclofenac | CXCL8 | 0.74 |
| fluticasone | NR3C1 | 0.74 |
| vemurafenib | MAP3K20 | 0.74 |
| everolimus | MTOR | 0.74 |
| nintedanib | MELK | 0.73 |
| fedratinib | JAK2 | 0.73 |
| abemaciclib | CAMK2G | 0.73 |
| nintedanib | KCNH2 | 0.73 |
| nilotinib | CA2 | 0.73 |
| dasatinib | TESK1 | 0.73 |
| risedronate | FDPS | 0.73 |
| erlotinib | SLK | 0.73 |
| cyclosporine | PPID | 0.73 |
| lovastatin | HMGCR | 0.73 |
| tofacitinib | DCLK3 | 0.73 |
| alectinib | EML4 | 0.73 |
| copanlisib | PIK3R1 | 0.73 |
| tivozanib | PEBP1 | 0.73 |
| teduglutide | GLP2R | 0.73 |
| sirolimus | FKBP1B | 0.72 |
| nintedanib | MAP3K7 | 0.72 |
| ibrutinib | CSK | 0.72 |
| dasatinib | SIK1 | 0.72 |
| romidepsin | NCOR1 | 0.72 |
| bosutinib | STK35 | 0.72 |
| erlotinib | BUB1 | 0.72 |
| midostaurin | PKN1 | 0.71 |
| vorinostat | NCOR2 | 0.71 |
| midostaurin | GRK7 | 0.71 |
| abemaciclib | CAMK2D | 0.71 |
| pentostatin | ADA | 0.71 |
| haloperidol | SIGMAR1 | 0.71 |
| dasatinib | YES1 | 0.71 |
| sirolimus | EIF4E | 0.71 |
| palbociclib | CCND1 | 0.71 |
| abemaciclib | GSK3A | 0.70 |
| idoxuridine | TK1 | 0.70 |
| nintedanib | PRPF4B | 0.70 |
| mafenide | CA12 | 0.70 |

| | | |
|---|---|---|
| mycophenolic | IMPDH1 | 0.70 |
| tamoxifen | EBP | 0.70 |
| midostaurin | GRK1 | 0.70 |
| ruxolitinib | HDAC6 | 0.70 |
| osimertinib | CARS1 | 0.70 |
| midostaurin | RIOK2 | 0.70 |
| clotrimazole | CYP19A1 | 0.70 |
| niraparib | PARP2 | 0.69 |
| pimozide | HTR7 | 0.69 |
| nintedanib | SRPK3 | 0.69 |
| tamoxifen | DHCR7 | 0.69 |
| dipyridamole | SLC29A1 | 0.69 |
| telmisartan | AGTR2 | 0.69 |
| fedratinib | DAPK1 | 0.69 |
| sunitinib | MAP4K1 | 0.69 |
| disulfiram | LOXL4 | 0.69 |
| dasatinib | EPHB3 | 0.69 |
| calcitriol | VDR | 0.69 |
| loperamide | OPRM1 | 0.69 |
| sunitinib | CHEK2 | 0.69 |
| prazosin | ADRA1D | 0.69 |
| losartan | AGTR1 | 0.69 |
| nintedanib | GRK4 | 0.69 |
| nintedanib | PIP4K2B | 0.69 |
| vandetanib | RIPK2 | 0.69 |
| selumetinib | MAP2K1 | 0.68 |
| sunitinib | PAK3 | 0.68 |
| methotrexate | DHFR | 0.68 |
| bosutinib | STK33 | 0.68 |
| fenofibrate | FABP1 | 0.68 |
| midostaurin | MARK1 | 0.68 |
| dasatinib | TXK | 0.68 |
| gefitinib | SBK1 | 0.68 |
| methotrexate | SLC19A1 | 0.68 |
| regorafenib | TEK | 0.68 |
| plerixafor | CCR2 | 0.68 |
| propranolol | ADRB2 | 0.68 |
| fedratinib | DAPK2 | 0.68 |
| nintedanib | RIOK1 | 0.68 |
| imatinib | SLC47A1 | 0.68 |
| citalopram | SLC6A4 | 0.68 |
| lonafarnib | FNTB | 0.68 |
| sorafenib | HIPK4 | 0.67 |
| pexidartinib | PRKCQ | 0.67 |
| ruxolitinib | CAMK2A | 0.67 |
| maraviroc | CCR5 | 0.67 |

| | | |
|---|---|---|
| indomethacin | NAPRT | 0.67 |
| dasatinib | TNNI3K | 0.67 |
| tretinoin | RARG | 0.67 |
| sincalide | CCKBR | 0.67 |
| oxymetazoline | HTR1B | 0.67 |
| sunitinib | STK39 | 0.67 |
| sirolimus | FKBP5 | 0.67 |
| rimegepant | CALCRL | 0.67 |
| aprepitant | TACR1 | 0.67 |
| etomidate | CYP11B1 | 0.67 |
| nintedanib | RIOK3 | 0.67 |
| pacritinib | STK11 | 0.67 |
| clonazepam | GABRA3 | 0.67 |
| abemaciclib | ADCK1 | 0.67 |
| vandetanib | EPHA6 | 0.66 |
| desmopressin | AVPR2 | 0.66 |
| digitoxin | ATP1A1 | 0.66 |
| digitoxin | ATP1B1 | 0.66 |
| digitoxin | ATP1A3 | 0.66 |
| digitoxin | ATP1B2 | 0.66 |
| digitoxin | ATP1A2 | 0.66 |
| digitoxin | ATP1B3 | 0.66 |
| digitoxin | FXYD2 | 0.66 |
| digitoxin | ATP1A4 | 0.66 |
| nifedipine | CACNA1F | 0.66 |
| nifedipine | CACNA1S | 0.66 |
| ruxolitinib | DCLK1 | 0.66 |
| neratinib | MINK1 | 0.66 |
| decitabine | DNMT1 | 0.66 |
| ceritinib | TNK1 | 0.66 |
| estradiol | GPER1 | 0.66 |
| cyclosporine | PPIA | 0.66 |
| abemaciclib | CLK1 | 0.66 |
| neratinib | PAK2 | 0.66 |
| estradiol | ESRRB | 0.66 |
| nintedanib | PIP5K1A | 0.66 |
| nisoldipine | CACNA1C | 0.66 |
| bexarotene | RXRB | 0.66 |
| propranolol | ADRB1 | 0.66 |
| tamoxifen | EBPL | 0.65 |
| dasatinib | SIK2 | 0.65 |
| sulfanilamide | CA13 | 0.65 |
| vorinostat | RCOR3 | 0.65 |
| sunitinib | ULK2 | 0.65 |
| trametinib | MAP2K7 | 0.65 |
| cyclosporine | FKBP4 | 0.65 |

| | | |
|---|---|---|
| dasatinib | EPHA1 | 0.65 |
| abemaciclib | GSK3B | 0.65 |
| selumetinib | SMC2 | 0.65 |
| ruxolitinib | CAMK1D | 0.65 |
| raloxifene | AOX1 | 0.65 |
| nintedanib | LRRK2 | 0.64 |
| ibrutinib | TEC | 0.64 |
| midostaurin | MAP3K9 | 0.64 |
| orlistat | LIPC | 0.64 |
| cangrelor | P2RY12 | 0.64 |
| bosutinib | BTK | 0.64 |
| nicotine | CHRNA2 | 0.64 |
| rosiglitazone | CISD1 | 0.64 |
| lenvatinib | KIF5B | 0.64 |
| lenvatinib | CCDC6 | 0.64 |
| prednisolone | SERPINA6 | 0.64 |
| imatinib | NULL | 0.64 |
| febuxostat | XDH | 0.64 |
| doxazosin | ADRA2A | 0.64 |
| ivacaftor | CFTR | 0.64 |
| palbociclib | PIP4K2A | 0.64 |
| liothyronine | THRA | 0.64 |
| selumetinib | SMC1A | 0.64 |
| sunitinib | CSNK1E | 0.64 |
| fluphenazine | DRD3 | 0.64 |
| erlotinib | STK10 | 0.64 |
| ponatinib | DDX1 | 0.64 |
| dasatinib | PKMYT1 | 0.64 |
| itraconazole | CYP3A4 | 0.63 |
| lemborexant | HCRTR2 | 0.63 |
| warfarin | VKORC1 | 0.63 |
| midostaurin | SRPK1 | 0.63 |
| tacrolimus | PPP3CA | 0.63 |
| liothyronine | THRB | 0.63 |
| paclitaxel | TUBB3 | 0.63 |
| bosutinib | MAP4K2 | 0.63 |
| dasatinib | TNK2 | 0.63 |
| imatinib | CA1 | 0.63 |
| imatinib | DDX3X | 0.63 |
| pyrilamine | HRH1 | 0.63 |
| ibrutinib | PTK6 | 0.63 |
| axitinib | CSF1R | 0.63 |
| ketoprofen | PTGS1 | 0.63 |
| pacritinib | TYK2 | 0.62 |
| ceritinib | TSSK1B | 0.62 |
| sunitinib | CSNK1D | 0.62 |

| | | |
|---|---|---|
| prazosin | ADRA1B | 0.62 |
| sunitinib | ITK | 0.62 |
| roflumilast | PDE4D | 0.62 |
| bosutinib | MAP4K3 | 0.62 |
| aripiprazole | HTR2B | 0.62 |
| nilotinib | ILK | 0.62 |
| cyclosporine | PPIB | 0.62 |
| lisinopril | ACE | 0.62 |
| niraparib | PARP3 | 0.62 |
| ambrisentan | EDNRA | 0.62 |
| nilotinib | CA9 | 0.62 |
| diclofenac | PTGS2 | 0.62 |
| carbachol | CHRM2 | 0.62 |
| butorphanol | OPRK1 | 0.62 |
| cobimetinib | ACTR3 | 0.62 |
| chlorpromazine | ADRA1A | 0.62 |
| sunitinib | HIPK2 | 0.61 |
| adapalene | RARB | 0.61 |
| sorafenib | RAF1 | 0.61 |
| doxazosin | ADRA2C | 0.61 |
| gefitinib | MKNK1 | 0.61 |
| tacrine | APP | 0.61 |
| neratinib | PAK1 | 0.61 |
| sirolimus | PDCD4 | 0.61 |
| mitoxantrone | ABCC1 | 0.61 |
| ruxolitinib | CAMK1 | 0.61 |
| dabrafenib | ARAF | 0.61 |
| nifedipine | CACNA1D | 0.61 |
| uridine | CDA | 0.61 |
| dasatinib | EPHB6 | 0.61 |
| paclitaxel | TUBB4A | 0.61 |
| paclitaxel | TUBB | 0.61 |
| paclitaxel | TUBA1B | 0.61 |
| paclitaxel | TUBA4A | 0.61 |
| paclitaxel | TUBB4B | 0.61 |
| paclitaxel | TUBB2A | 0.61 |
| paclitaxel | TUBB8 | 0.61 |
| paclitaxel | TUBA3E | 0.61 |
| paclitaxel | TUBA1A | 0.61 |
| paclitaxel | TUBA1C | 0.61 |
| paclitaxel | TUBB6 | 0.61 |
| paclitaxel | TUBB2B | 0.61 |
| paclitaxel | TUBB1 | 0.61 |
| fedratinib | SNRK | 0.61 |
| midostaurin | NUAK1 | 0.61 |
| methotrexate | SLC46A1 | 0.61 |

| | | |
|---|---|---|
| abemaciclib | CSNK2A2 | 0.60 |
| ceritinib | CAMK4 | 0.60 |
| midostaurin | TBK1 | 0.60 |
| pregabalin | CACNA2D1 | 0.60 |
| afamelanotide | MC1R | 0.60 |
| estradiol | SULT1E1 | 0.60 |
| lansoprazole | MAPT | 0.60 |
| abiraterone | CYP21A2 | 0.60 |
| tazemetostat | AEBP2 | 0.60 |
| bosutinib | TNIK | 0.60 |
| midostaurin | MARK3 | 0.60 |
| disulfiram | LOXL3 | 0.60 |
| nintedanib | SRPK2 | 0.60 |
| desipramine | SLC6A2 | 0.60 |
| iloprost | PTGIR | 0.60 |
| vardenafil | PDE5A | 0.60 |
| dasatinib | MYT1 | 0.60 |
| clotrimazole | TBXAS1 | 0.60 |
| trametinib | MAP2K2 | 0.60 |
| efavirenz | CYP2B6 | 0.59 |
| osilodrostat | CYP11B2 | 0.59 |
| pazopanib | STK36 | 0.59 |
| sunitinib | TLK2 | 0.59 |
| copanlisib | PIK3CG | 0.59 |
| rosiglitazone | PPARG | 0.59 |
| nintedanib | STK4 | 0.59 |
| sincalide | CCKAR | 0.59 |
| vandetanib | TYRO3 | 0.59 |
| midostaurin | CAMK2B | 0.59 |
| abemaciclib | IRAK1 | 0.59 |
| dutasteride | SRD5A1 | 0.59 |
| cytarabine | MDM2 | 0.59 |
| tretinoin | RARA | 0.59 |
| ramelteon | MTNR1B | 0.59 |
| abemaciclib | CILK1 | 0.59 |
| midostaurin | IKBKE | 0.59 |
| midostaurin | MARK2 | 0.59 |
| ramelteon | MTNR1A | 0.59 |
| ceritinib | INSR | 0.59 |
| tivozanib | FGF2 | 0.59 |
| oxytocin | OXTR | 0.59 |
| estrone | SHBG | 0.58 |
| abiraterone | CYP17A1 | 0.58 |
| olaparib | PARP1 | 0.58 |
| sorafenib | CCNC | 0.58 |
| vismodegib | SHH | 0.58 |

| | | |
|---|---|---|
| disulfiram | LOX | 0.58 |
| axitinib | TIE1 | 0.58 |
| dasatinib | SRMS | 0.58 |
| neratinib | EIF2AK4 | 0.58 |
| tacrolimus | SF3B3 | 0.58 |
| tirofiban | ITGA2B | 0.58 |
| colchicine | SLC22A3 | 0.58 |
| abemaciclib | CDK16 | 0.58 |
| bexarotene | NR4A2 | 0.58 |
| glucagon | GCGR | 0.58 |
| nintedanib | MERTK | 0.58 |
| migalastat | GLA | 0.58 |
| mycophenolic | IMPDH2 | 0.58 |
| dabrafenib | ERN2 | 0.58 |
| ribociclib | CDK9 | 0.58 |
| dabigatran | F2 | 0.58 |
| sunitinib | PHKG2 | 0.58 |
| diazepam | GABRP | 0.57 |
| diazepam | GABRD | 0.57 |
| diazepam | GABRA4 | 0.57 |
| diazepam | GABRE | 0.57 |
| diazepam | GABRG1 | 0.57 |
| diazepam | GABRG3 | 0.57 |
| diazepam | GABRQ | 0.57 |
| lenalidomide | IKZF1 | 0.57 |
| trametinib | ABCB1 | 0.57 |
| propranolol | ADRB3 | 0.57 |
| saxagliptin | DPP4 | 0.57 |
| diazepam | GABRB1 | 0.57 |
| sunitinib | AAK1 | 0.57 |
| telithromycin | CYP3A5 | 0.57 |
| telithromycin | CYP3A7 | 0.57 |
| telithromycin | CYP3A43 | 0.57 |
| lemborexant | HCRTR1 | 0.57 |
| losartan | SLCO1B1 | 0.57 |
| cabozantinib | AXL | 0.57 |
| apixaban | F10 | 0.57 |
| sunitinib | PRKAA1 | 0.57 |
| sunitinib | CSNK1A1L | 0.57 |
| dinoprostone | PTGER4 | 0.57 |
| crizotinib | MLKL | 0.57 |
| lenvatinib | MAPKAPK 2 | 0.57 |
| midostaurin | CAMKK2 | 0.57 |
| ceftriaxone | SLC22A6 | 0.57 |
| lenalidomide | IKZF3 | 0.57 |

| | | |
|---|---|---|
| reserpine | SLC18A2 | 0.57 |
| nintedanib | INSRR | 0.57 |
| calcitriol | RXRA | 0.57 |
| ibrutinib | RIPK3 | 0.57 |
| crizotinib | SBK3 | 0.57 |
| mupirocin | IARS1 | 0.57 |
| chlorpromazine | HTR2C | 0.56 |
| naproxen | AKR1C3 | 0.56 |
| sunitinib | OXSR1 | 0.56 |
| prednisolone | GLUL | 0.56 |
| granisetron | HTR3B | 0.56 |
| clotrimazole | NR1I3 | 0.56 |
| haloperidol | TMEM97 | 0.56 |
| sunitinib | BMP2K | 0.56 |
| neratinib | MAP3K12 | 0.56 |
| disulfiram | LOXL2 | 0.56 |
| erlotinib | SLCO2B1 | 0.56 |
| midostaurin | PRKG2 | 0.56 |
| afamelanotide | MC3R | 0.56 |
| abemaciclib | PIM1 | 0.56 |
| clotrimazole | CYP51A1 | 0.56 |
| roflumilast | PDE4B | 0.56 |
| chlorpromazine | HTR2A | 0.56 |
| fostamatinib | NEK9 | 0.56 |
| dabrafenib | PRKD2 | 0.56 |
| prednisolone | NR3C2 | 0.56 |
| axitinib | AURKB | 0.55 |
| docetaxel | GHRHR | 0.55 |
| sunitinib | STK17A | 0.55 |
| adenosine | ADORA1 | 0.55 |
| lonafarnib | HRAS | 0.55 |
| afamelanotide | MC5R | 0.55 |
| clozapine | H1-0 | 0.55 |
| midostaurin | SYK | 0.55 |
| bosutinib | MAP3K3 | 0.55 |
| imatinib | CA3 | 0.55 |
| sumatriptan | HTR1D | 0.55 |
| orlistat | LIPG | 0.55 |
| bosutinib | WEE2 | 0.55 |
| afamelanotide | MC4R | 0.55 |
| nintedanib | FGFR1 | 0.55 |
| imatinib | NQO2 | 0.55 |
| sunitinib | MYLK2 | 0.55 |
| abemaciclib | TAOK1 | 0.55 |
| bortezomib | PSMD11 | 0.55 |
| bortezomib | PSMD12 | 0.55 |

| | | |
|---|---|---|
| bortezomib | PSMD14 | 0.55 |
| bortezomib | PSMD3 | 0.55 |
| bortezomib | PSMC3 | 0.55 |
| bortezomib | PSMC2 | 0.55 |
| bortezomib | PSMD8 | 0.55 |
| bortezomib | PSMD7 | 0.55 |
| bortezomib | PSMD4 | 0.55 |
| bortezomib | SEM1 | 0.55 |
| bortezomib | PSMC1 | 0.55 |
| bortezomib | PSMC5 | 0.55 |
| bortezomib | PSMC6 | 0.55 |
| bortezomib | PSMD2 | 0.55 |
| bortezomib | PSMD6 | 0.55 |
| bortezomib | ADRM1 | 0.55 |
| bortezomib | PSMD1 | 0.55 |
| bortezomib | PSMD13 | 0.55 |
| abemaciclib | CLK4 | 0.55 |
| histamine | HRH4 | 0.55 |
| midostaurin | BRSK2 | 0.55 |
| sunitinib | HIPK3 | 0.55 |
| levorphanol | OPRD1 | 0.55 |
| miglustat | GAA | 0.55 |
| sunitinib | MAP4K4 | 0.54 |
| crizotinib | LTK | 0.54 |
| pemetrexed | FOLR1 | 0.54 |
| fedratinib | STK16 | 0.54 |
| nicotine | CHRNA6 | 0.54 |
| bortezomib | PSMB11 | 0.54 |
| bortezomib | PSMA7 | 0.54 |
| bortezomib | PSMA1 | 0.54 |
| bortezomib | PSMA2 | 0.54 |
| bortezomib | PSMA3 | 0.54 |
| bortezomib | PSMA4 | 0.54 |
| bortezomib | PSMA5 | 0.54 |
| bortezomib | PSMB4 | 0.54 |
| bortezomib | PSMB6 | 0.54 |
| bortezomib | PSMB3 | 0.54 |
| bortezomib | PSMA6 | 0.54 |
| bortezomib | PSMA8 | 0.54 |
| bortezomib | PSMB7 | 0.54 |
| nintedanib | MAP2K5 | 0.54 |
| clotrimazole | CYP2C9 | 0.54 |
| granisetron | HTR3A | 0.54 |
| sunitinib | STK3 | 0.54 |
| midostaurin | CAMKK1 | 0.54 |
| sunitinib | SGK3 | 0.54 |

| | | |
|---|---|---|
| ruxolitinib | ANKK1 | 0.54 |
| fedratinib | PTK2 | 0.54 |
| lifitegrast | ICAM1 | 0.54 |
| fulvestrant | ESRRA | 0.54 |
| axitinib | AURKA | 0.54 |
| carbachol | CHRM5 | 0.54 |
| tadalafil | PDE11A | 0.54 |
| pazopanib | RIPK1 | 0.54 |
| roflumilast | PDE4C | 0.54 |
| histamine | CA14 | 0.54 |
| phenelzine | AOC3 | 0.53 |
| dronabinol | CNR1 | 0.53 |
| vasopressin | AVPR1A | 0.53 |
| lenvatinib | MAPK14 | 0.53 |
| neratinib | MAP3K4 | 0.53 |
| abemaciclib | DYRK1A | 0.53 |
| copanlisib | PIK3CB | 0.53 |
| desmopressin | AVPR1B | 0.53 |
| roflumilast | PDE4A | 0.53 |
| methoxsalen | CYP1A2 | 0.53 |
| glyburide | ABCC8 | 0.53 |
| methotrexate | HMGB1 | 0.53 |
| midostaurin | IRAK3 | 0.53 |
| dasatinib | TGFBR1 | 0.53 |
| neratinib | MAP3K13 | 0.53 |
| zafirlukast | CYSLTR2 | 0.53 |
| panobinostat | HDAC3 | 0.53 |
| dasatinib | ACVR2A | 0.53 |
| raltegravir | CCR1 | 0.53 |
| sunitinib | MAST1 | 0.53 |
| gilteritinib | ULK3 | 0.53 |
| abemaciclib | CCNA2 | 0.53 |
| levothyroxine | SLCO1C1 | 0.53 |
| panobinostat | HDAC1 | 0.53 |
| bosutinib | MAP3K2 | 0.53 |
| clotrimazole | KCNN4 | 0.53 |
| dronabinol | CNR2 | 0.53 |
| fluoxetine | CYP2C19 | 0.53 |
| pacritinib | ACVR1 | 0.53 |
| tazemetostat | EZH2 | 0.53 |
| bosutinib | DSTYK | 0.53 |
| levodopa | CA5B | 0.53 |
| tucatinib | MYH14 | 0.53 |
| sunitinib | PRKD1 | 0.52 |
| midostaurin | HUNK | 0.52 |
| sunitinib | CLK2 | 0.52 |

| | | |
|---|---|---|
| midostaurin | TTK | 0.52 |
| indomethacin | PTGER2 | 0.52 |
| pioglitazone | ABCB11 | 0.52 |
| bortezomib | NFKB1 | 0.52 |
| bortezomib | NFKB2 | 0.52 |
| axitinib | CDKL2 | 0.52 |
| pimozide | SCN11A | 0.52 |
| capsaicin | TRPV1 | 0.52 |
| ranitidine | ATP4A | 0.52 |
| ranitidine | ATP4B | 0.52 |
| pazopanib | MAP3K11 | 0.52 |
| adenosine | ADORA3 | 0.52 |
| bosutinib | HIPK1 | 0.52 |
| nintedanib | CAMK1G | 0.52 |
| mesalamine | TST | 0.52 |
| selegiline | MAOB | 0.52 |
| dabrafenib | CDK2 | 0.52 |
| sunitinib | RPS6KA3 | 0.52 |
| fedratinib | COQ8B | 0.52 |
| sunitinib | RPS6KA1 | 0.52 |
| istradefylline | ADORA2A | 0.52 |
| pazopanib | PI4KB | 0.52 |
| sunitinib | EIF2AK2 | 0.52 |
| bosutinib | NEK11 | 0.52 |
| sunitinib | STK38 | 0.52 |
| fedratinib | STK17B | 0.52 |
| ponatinib | FGFR4 | 0.52 |
| midostaurin | PRKCH | 0.51 |
| alprostadil | SLCO3A1 | 0.51 |
| ruxolitinib | ROCK1 | 0.51 |
| orlistat | ABHD16A | 0.51 |
| fedratinib | NEK6 | 0.51 |
| tiotropium | CHRM3 | 0.51 |
| ozanimod | S1PR1 | 0.51 |
| captopril | REN | 0.51 |
| midostaurin | DYRK1B | 0.51 |
| alectinib | CSNK1G2 | 0.51 |
| tegaserod | HTR4 | 0.51 |
| sunitinib | PHKG1 | 0.51 |
| estradiol | HSD17B1 | 0.51 |
| tacrine | HNMT | 0.51 |
| fedratinib | MAPK7 | 0.51 |
| bosutinib | DMPK | 0.51 |
| nintedanib | PRKAA2 | 0.51 |
| haloperidol | DRD5 | 0.51 |
| tazemetostat | SUZ12 | 0.51 |

| | | |
|---|---|---|
| nintedanib | LATS2 | 0.51 |
| dasatinib | LIMK1 | 0.51 |
| ribociclib | CCNT1 | 0.51 |
| pemetrexed | GART | 0.51 |
| sunitinib | NIM1K | 0.51 |
| abemaciclib | PRKCA | 0.51 |
| ruxolitinib | MKNK2 | 0.50 |
| ceritinib | FER | 0.50 |
| cefotaxime | SLC22A8 | 0.50 |
| crizotinib | MST1R | 0.50 |
| sunitinib | JAK1 | 0.50 |
| erlotinib | MAPK4 | 0.50 |
| sunitinib | TLK1 | 0.50 |
| panobinostat | HDAC2 | 0.50 |
| tazemetostat | RBBP4 | 0.50 |
| sunitinib | CDK14 | 0.50 |
| bosentan | EDNRB | 0.50 |
| dasatinib | LIMK2 | 0.50 |
| pentamidine | PTP4A2 | 0.50 |
| pentamidine | PTP4A1 | 0.50 |
| midostaurin | PRKCE | 0.50 |
| dinoprost | PTGFR | 0.50 |
| dapagliflozin | SLC5A2 | 0.50 |
| sunitinib | CDK7 | 0.50 |
| dasatinib | SIK3 | 0.50 |
| bosutinib | WEE1 | 0.50 |
| danazol | CYP2J2 | 0.50 |
| sunitinib | DYRK2 | 0.50 |
| granisetron | HTR3E | 0.50 |
| granisetron | HTR3D | 0.50 |
| granisetron | HTR3C | 0.50 |
| nintedanib | CDK18 | 0.50 |
| thioridazine | CHRM1 | 0.50 |
| nicotine | CHRNB3 | 0.49 |
| abemaciclib | PRKCD | 0.49 |
| encorafenib | PCYT1A | 0.49 |
| midostaurin | PIM3 | 0.49 |
| lenvatinib | CA5A | 0.49 |
| midostaurin | PDPK1 | 0.49 |
| midostaurin | PIK3C2G | 0.49 |
| sunitinib | CSNK2A1 | 0.49 |
| sunitinib | CHEK1 | 0.49 |
| lanreotide | SSTR2 | 0.49 |
| risdiplam | SMN1 | 0.49 |
| nintedanib | MUSK | 0.49 |
| sunitinib | STK26 | 0.49 |

| | | |
|---|---|---|
| sulfanilamide | CA7 | 0.49 |
| diazepam | GABRA5 | 0.49 |
| vilazodone | HTR1A | 0.49 |
| voriconazole | CYP46A1 | 0.49 |
| haloperidol | DRD1 | 0.49 |
| methotrexate | FOLR2 | 0.49 |
| alectinib | ACOX1 | 0.49 |
| tacrine | CHRNE | 0.49 |
| sunitinib | CHUK | 0.49 |
| indomethacin | PTGDR2 | 0.49 |
| tamoxifen | ESRRG | 0.49 |
| lenvatinib | NCOA4 | 0.48 |
| ponatinib | MYLK | 0.48 |
| midostaurin | PRKX | 0.48 |
| migalastat | SI | 0.48 |
| regorafenib | MAP3K1 | 0.48 |
| ixazomib | PSMB5 | 0.48 |
| zalcitabine | CXCR4 | 0.48 |
| vardenafil | PDE6D | 0.48 |
| vardenafil | PDE6G | 0.48 |
| vardenafil | PDE6B | 0.48 |
| vardenafil | PDE6H | 0.48 |
| chlorpromazine | HTR6 | 0.48 |
| diazepam | GABRA2 | 0.48 |
| sunitinib | BMPR2 | 0.48 |
| lenalidomide | CRBN | 0.48 |
| dinoprostone | SLC22A2 | 0.48 |
| sunitinib | DLK1 | 0.48 |
| ciprofloxacin | TOP2B | 0.48 |
| midostaurin | TNKS2 | 0.48 |
| telaprevir | CELA1 | 0.48 |
| dabrafenib | PRKD3 | 0.48 |
| sunitinib | LATS1 | 0.48 |
| sunitinib | PAK5 | 0.48 |
| tacrine | BCHE | 0.47 |
| erythromycin | MLNR | 0.47 |
| vandetanib | ACVRL1 | 0.47 |
| lapatinib | PIK3C2B | 0.47 |
| tazemetostat | EED | 0.47 |
| belinostat | HDAC9 | 0.47 |
| olaparib | PARP4 | 0.47 |
| dasatinib | NLK | 0.47 |
| panobinostat | HDAC10 | 0.47 |
| orlistat | LPL | 0.47 |
| dinoprostone | PTGER3 | 0.47 |
| dabrafenib | CDK17 | 0.47 |

| | | |
|---|---|---|
| lifitegrast | ITGB2 | 0.47 |
| chlorpromazine | ADRA2B | 0.47 |
| gefitinib | MAPK6 | 0.47 |
| zileuton | ALOX5 | 0.47 |
| sunitinib | RPS6KA2 | 0.47 |
| fedratinib | MAPK9 | 0.47 |
| ixazomib | PSMB10 | 0.47 |
| ceritinib | IGF1R | 0.47 |
| ondansetron | SLC47A2 | 0.47 |
| sorafenib | CDK19 | 0.47 |
| crizotinib | CASK | 0.46 |
| ruxolitinib | NUAK2 | 0.46 |
| neostigmine | ACHE | 0.46 |
| isoproterenol | TAS2R41 | 0.46 |
| isoproterenol | TAS2R43 | 0.46 |
| isoproterenol | TAS2R60 | 0.46 |
| isoproterenol | TAS2R14 | 0.46 |
| isoproterenol | TAS2R13 | 0.46 |
| isoproterenol | TAS2R9 | 0.46 |
| isoproterenol | TAS2R4 | 0.46 |
| isoproterenol | TAS2R1 | 0.46 |
| alprostadil | SLCO2A1 | 0.46 |
| gilteritinib | ETV6 | 0.46 |
| pimozide | SCN5A | 0.46 |
| dasatinib | ACVR1B | 0.46 |
| neratinib | ADCK5 | 0.46 |
| gefitinib | ABCC4 | 0.46 |
| orlistat | PNLIP | 0.46 |
| sunitinib | CSNK1G3 | 0.46 |
| abemaciclib | CCNE1 | 0.46 |
| cromolyn | GPR35 | 0.46 |
| amiloride | SCNN1A | 0.46 |
| encorafenib | PGRMC1 | 0.46 |
| paroxetine | MPO | 0.46 |
| paclitaxel | ITGB3 | 0.46 |
| flumazenil | GABRG2 | 0.46 |
| flumazenil | GABRB3 | 0.46 |
| diazepam | GABRA1 | 0.46 |
| doxorubicin | SF1 | 0.46 |
| memantine | GRIN2D | 0.46 |
| memantine | GRIN2C | 0.46 |
| histamine | HRH3 | 0.46 |
| dasatinib | ACVR2B | 0.46 |
| niclosamide | STAT3 | 0.46 |
| guanfacine | NISCH | 0.45 |
| tiagabine | SLC6A1 | 0.45 |

| | | |
|---|---|---|
| pimozide | SCN4A | 0.45 |
| sertraline | SLC6A3 | 0.45 |
| imatinib | CA6 | 0.45 |
| ponatinib | PTK2B | 0.45 |
| topiramate | CA11 | 0.45 |
| dapagliflozin | SLC5A4 | 0.45 |
| fedratinib | MATK | 0.45 |
| cilostazol | PDE3B | 0.45 |
| cilostazol | PDE3A | 0.45 |
| sorafenib | CDK8 | 0.45 |
| bosutinib | CLK3 | 0.45 |
| nilotinib | CA4 | 0.45 |
| ruxolitinib | DCLK2 | 0.45 |
| dopamine | DRD4 | 0.45 |
| bosutinib | VRK2 | 0.45 |
| vismodegib | SMO | 0.45 |
| fedratinib | NEK5 | 0.45 |
| paclitaxel | ITGAV | 0.45 |
| sorafenib | CDKL3 | 0.45 |
| ruxolitinib | PLK3 | 0.44 |
| orlistat | DAGLA | 0.44 |
| ruxolitinib | PLK1 | 0.44 |
| memantine | GRIN3B | 0.44 |
| memantine | GRIN3A | 0.44 |
| panobinostat | HDAC11 | 0.44 |
| lomitapide | MTTP | 0.44 |
| abemaciclib | CIT | 0.44 |
| fedratinib | NEK4 | 0.44 |
| fedratinib | BMPR1B | 0.44 |
| idelalisib | PIK3R2 | 0.44 |
| fedratinib | IKBKB | 0.44 |
| saxagliptin | DPP9 | 0.44 |
| bortezomib | PSMB9 | 0.44 |
| tolvaptan | CRHR1 | 0.44 |
| disulfiram | EHMT2 | 0.44 |
| dabrafenib | NEK1 | 0.44 |
| captopril | ACE2 | 0.44 |
| lindane | GABRA6 | 0.44 |
| belzutifan | EPAS1 | 0.43 |
| hexachlorophene | HSPD1 | 0.43 |
| hexachlorophene | HSPE1 | 0.43 |
| belinostat | HDAC7 | 0.43 |
| pimozide | SCN1A | 0.43 |
| panobinostat | HDAC8 | 0.43 |
| pemetrexed | ATIC | 0.43 |
| lifitegrast | ITGAL | 0.43 |

| | | |
|---|---|---|
| dabrafenib | ULK1 | 0.43 |
| glyburide | KCNJ11 | 0.43 |
| orlistat | ABHD6 | 0.43 |
| sunitinib | CSNK1A1 | 0.43 |
| eltrombopag | MPL | 0.43 |
| sunitinib | CSNK1G1 | 0.43 |
| gilteritinib | CYC1 | 0.43 |
| bortezomib | PSMB8 | 0.43 |
| dexamethasone | PLA2G1B | 0.42 |
| atorvastatin | SLCO1B3 | 0.42 |
| probucol | HPN | 0.42 |
| cangrelor | GPR17 | 0.42 |
| rosiglitazone | CISD2 | 0.42 |
| acetylcholine | CHRM4 | 0.42 |
| nicotine | CHRNA1 | 0.42 |
| doxorubicin | HIF1A | 0.42 |
| isoproterenol | TAS2R46 | 0.42 |
| vardenafil | PDE6A | 0.42 |
| cephalothin | SLC22A11 | 0.42 |
| baclofen | GABBR1 | 0.42 |
| isotretinoin | OR51E2 | 0.42 |
| bosutinib | PLK2 | 0.42 |
| nicotine | CHRNG | 0.42 |
| nicotine | CHRNB1 | 0.42 |
| axitinib | MYO3A | 0.42 |
| zileuton | ALOX15 | 0.42 |
| lenalidomide | IL1B | 0.42 |
| baclofen | GABBR2 | 0.42 |
| bosutinib | MYO3B | 0.42 |
| pimozide | SCN3A | 0.42 |
| nicotine | CHRND | 0.42 |
| isoproterenol | TAS2R31 | 0.42 |
| ezogabine | KCNQ5 | 0.41 |
| adenosine | SLC28A1 | 0.41 |
| methoxsalen | CYP2A13 | 0.41 |
| sildenafil | PDE6C | 0.41 |
| glyburide | ABCC9 | 0.41 |
| sunitinib | GRK6 | 0.41 |
| fedratinib | TGFBR2 | 0.41 |
| pimozide | SCN2A | 0.41 |
| panobinostat | HDAC4 | 0.41 |
| disulfiram | MGLL | 0.41 |
| ranitidine | HRH2 | 0.41 |
| calcitriol | GC | 0.41 |
| diethylstilbestrol | ABCG2 | 0.41 |
| adenosine | AHCY | 0.41 |

| | | |
|---|---|---|
| varenicline | CHRNB2 | 0.41 |
| belinostat | HDAC5 | 0.41 |
| erlotinib | ACAD11 | 0.41 |
| tolcapone | TTR | 0.41 |
| pentamidine | AOC1 | 0.41 |
| crizotinib | EPHA7 | 0.41 |
| varenicline | CHRNA4 | 0.41 |
| pimozide | CACNA1G | 0.41 |
| baricitinib | EIF2AK1 | 0.40 |
| loratadine | SLC6A15 | 0.40 |
| crizotinib | CDK11A | 0.40 |
| abemaciclib | CDK5R1 | 0.40 |
| pimozide | SCN8A | 0.40 |
| gilteritinib | TUFM | 0.40 |
| fedratinib | BRD4 | 0.40 |
| fedratinib | BRDT | 0.40 |
| dabrafenib | CDK1 | 0.40 |
| dinoprostone | PTGER1 | 0.40 |
| hydroxychloroquine | TLR9 | 0.40 |
| isoproterenol | TAS2R10 | 0.40 |
| bortezomib | RELA | 0.40 |
| sunitinib | RPS6KA5 | 0.39 |
| ergotamine | HTR5A | 0.39 |
| indomethacin | COX2 | 0.39 |
| prednisolone | TNF | 0.39 |
| dabigatran | PRSS2 | 0.39 |
| dabigatran | PRSS3 | 0.39 |
| entacapone | COMT | 0.39 |
| bortezomib | PSMC4 | 0.39 |
| orlistat | ABHD12 | 0.39 |
| baricitinib | SEPTIN9 | 0.39 |
| midostaurin | BRSK1 | 0.39 |
| ibuprofen | CXCR1 | 0.39 |
| crizotinib | CDK11B | 0.39 |
| miglustat | GBA2 | 0.39 |
| isoproterenol | TAS2R16 | 0.39 |
| baricitinib | ROCK2 | 0.39 |
| sunitinib | RPS6KA4 | 0.39 |
| telaprevir | CMA1 | 0.39 |
| cabozantinib | FECH | 0.39 |
| clotrimazole | CYP8B1 | 0.38 |
| dinoprost | SLC22A7 | 0.38 |
| sulindac | C5 | 0.38 |
| primaquine | SHMT2 | 0.38 |
| digitoxin | SLCO4C1 | 0.38 |

| | | |
|---|---|---|
| methylergonovine | HTR1F | 0.38 |
| chlorhexidine | SLC22A1 | 0.38 |
| dipyridamole | PDE7B | 0.38 |
| gilteritinib | DHCR24 | 0.38 |
| dipyridamole | PDE8A | 0.38 |
| pemetrexed | FPGS | 0.38 |
| ezogabine | KCNQ2 | 0.38 |
| memantine | GRIN2A | 0.38 |
| ibrutinib | PRXL2A | 0.38 |
| dabigatran | PRSS1 | 0.38 |
| amlodipine | KCNK2 | 0.38 |
| bexarotene | NR1H3 | 0.38 |
| venetoclax | BCL2 | 0.38 |
| adenosine | SLC28A3 | 0.38 |
| gilteritinib | RAN | 0.38 |
| pimozide | SCN7A | 0.38 |
| gilteritinib | SLC25A5 | 0.37 |
| niraparib | PARP12 | 0.37 |
| sunitinib | MAP3K15 | 0.37 |
| pimozide | SCN9A | 0.37 |
| zileuton | PTGES | 0.37 |
| prednisolone | GPBAR1 | 0.37 |
| sunitinib | IRAK4 | 0.37 |
| boceprevir | CTSV | 0.37 |
| tazemetostat | EZH1 | 0.37 |
| zafirlukast | MAPK1 | 0.37 |
| pimozide | SCN10A | 0.37 |
| baricitinib | PKN2 | 0.37 |
| rucaparib | TNKS | 0.37 |
| lanreotide | SSTR5 | 0.37 |
| buprenorphine | OPRL1 | 0.37 |
| vorinostat | LTA4H | 0.37 |
| fedratinib | MAPK8 | 0.36 |
| disulfiram | CASP1 | 0.36 |
| copanlisib | RAB27A | 0.36 |
| dasatinib | CDC42BPG | 0.36 |
| disulfiram | EHMT1 | 0.36 |
| flumazenil | GABRB2 | 0.36 |
| lenvatinib | ERN1 | 0.36 |
| memantine | GRIN2B | 0.36 |
| pentamidine | GRIN1 | 0.36 |
| bosutinib | CDK15 | 0.36 |
| midostaurin | ZAP70 | 0.36 |
| atazanavir | UGT1A1 | 0.36 |
| pazopanib | MAP3K10 | 0.36 |
| sulindac | GLRA1 | 0.36 |

| | | |
|---|---|---|
| sunitinib | PRKAG2 | 0.35 |
| clotrimazole | CYP2C8 | 0.35 |
| siponimod | S1PR5 | 0.35 |
| exenatide | GLP1R | 0.35 |
| iloprost | PTGDR | 0.35 |
| tretinoin | RORA | 0.35 |
| ezogabine | KCNQ3 | 0.35 |
| methoxsalen | CYP2A6 | 0.35 |
| chloroquine | PRNP | 0.35 |
| olaparib | PARP6 | 0.35 |
| chloroxine | ALOX12 | 0.35 |
| levobupivacaine | CYP2D6 | 0.35 |
| abemaciclib | CCNB1 | 0.35 |
| venetoclax | BCL2L2 | 0.35 |
| linagliptin | FAP | 0.35 |
| abemaciclib | PRKCB | 0.35 |
| dicumarol | NQO1 | 0.35 |
| dipyridamole | PRUNE1 | 0.35 |
| midostaurin | MAP2K4 | 0.34 |
| bosutinib | STK32A | 0.34 |
| lenvatinib | MAP3K6 | 0.34 |
| axitinib | MYLK3 | 0.34 |
| isoproterenol | GRK2 | 0.34 |
| saxagliptin | DPP8 | 0.34 |
| ibuprofen | CXCR2 | 0.34 |
| cabozantinib | PIP4K2C | 0.34 |
| tretinoin | PIN1 | 0.34 |
| methylergonovine | HTR1E | 0.34 |
| afatinib | ADK | 0.34 |
| lorlatinib | FES | 0.34 |
| gilteritinib | BMPR1A | 0.33 |
| gemcitabine | DCK | 0.33 |
| clofarabine | PDE2A | 0.33 |
| haloprogin | GALE | 0.33 |
| leuprolide | GNRHR | 0.33 |
| miconazole | IDO1 | 0.33 |
| nicotine | CHRNA7 | 0.33 |
| cannabidiol | CYP1A1 | 0.33 |
| venetoclax | MCL1 | 0.33 |
| palbociclib | CCNA1 | 0.33 |
| adenosine | SLC28A2 | 0.33 |
| icatibant | BDKRB2 | 0.33 |
| raloxifene | PLD2 | 0.33 |
| disulfiram | TRPA1 | 0.32 |
| gilteritinib | NEK2 | 0.32 |

| | | |
|---|---|---|
| dextroamphetamine | TAAR1 | 0.32 |
| pentamidine | MAOA | 0.32 |
| boceprevir | CTSK | 0.32 |
| zafirlukast | SLC10A1 | 0.32 |
| pazopanib | PIP5K1C | 0.32 |
| lanreotide | SSTR3 | 0.32 |
| deferoxamine | DOHH | 0.32 |
| pyrvinium | SYNJ1 | 0.32 |
| digoxin | RORC | 0.32 |
| chenodiol | SLC10A2 | 0.32 |
| bortezomib | CTRB1 | 0.32 |
| nimodipine | NR1I2 | 0.32 |
| bexarotene | NR1H2 | 0.32 |
| liothyronine | PCNA | 0.32 |
| disulfiram | GSDMD | 0.32 |
| ponesimod | S1PR3 | 0.32 |
| dipyridamole | PDE7A | 0.31 |
| sunitinib | PRKAG1 | 0.31 |
| vorapaxar | F2R | 0.31 |
| dactinomycin | GRB2 | 0.31 |
| nicotine | CHRNA3 | 0.31 |
| fulvestrant | NR1H4 | 0.31 |
| nicotine | CHRNB4 | 0.31 |
| telotristat | TPH1 | 0.31 |
| sulindac | AKR1B1 | 0.31 |
| teriflunomide | DHODH | 0.31 |
| fexofenadine | SLCO1A2 | 0.31 |
| sildenafil | PDE1C | 0.31 |
| isoproterenol | TAS2R38 | 0.31 |
| isoproterenol | TAS2R39 | 0.31 |
| isoproterenol | TAS2R40 | 0.31 |
| isoproterenol | TAS2R30 | 0.31 |
| isoproterenol | TAS2R19 | 0.31 |
| isoproterenol | TAS2R20 | 0.31 |
| isoproterenol | TAS2R50 | 0.31 |
| isoproterenol | TAS2R8 | 0.31 |
| isoproterenol | TAS2R7 | 0.31 |
| isoproterenol | TAS2R5 | 0.31 |
| isoproterenol | TAS2R3 | 0.31 |
| bexarotene | CYP26B1 | 0.31 |
| carfilzomib | PSMB1 | 0.30 |
| carfilzomib | GSTO1 | 0.30 |
| sunitinib | ERCC2 | 0.30 |
| alitretinoin | NR4A1 | 0.30 |
| mercaptopurine | HPRT1 | 0.30 |

| | | |
|---|---|---|
| imipramine | SMPD1 | 0.30 |
| abemaciclib | TAOK3 | 0.30 |
| erlotinib | DCTPP1 | 0.30 |
| pemetrexed | TYMS | 0.30 |
| sildenafil | PDE1A | 0.30 |
| trifluoperazine | CALM1 | 0.30 |
| donepezil | BACE1 | 0.30 |
| belzutifan | VEGFA | 0.30 |
| ponatinib | MAPK11 | 0.30 |
| miltefosine | AKT1 | 0.30 |
| dinoprostone | HPGDS | 0.30 |
| ubrogepant | RAMP1 | 0.30 |
| carfilzomib | PSMB2 | 0.29 |
| sunitinib | PAK6 | 0.29 |
| topotecan | TOP1 | 0.29 |
| fedratinib | RPS6KB1 | 0.29 |
| bortezomib | CTSG | 0.29 |
| ruxolitinib | STRADA | 0.29 |
| neratinib | STK32B | 0.29 |
| menadione | CDC25A | 0.29 |
| migalastat | MGAM | 0.29 |
| haloperidol | TAC1 | 0.29 |
| hexachlorophene | HSPA5 | 0.29 |
| gilteritinib | MAPK15 | 0.29 |
| diclofenac | AKR1B10 | 0.29 |
| zanamivir | NEU3 | 0.29 |
| estradiol | SULT1A1 | 0.28 |
| ruxolitinib | TAOK2 | 0.28 |
| idelalisib | PIK3C3 | 0.28 |
| ruxolitinib | NEK7 | 0.28 |
| pyrimethamine | HEXB | 0.28 |
| orlistat | PLA2G7 | 0.28 |
| alectinib | ACOX3 | 0.28 |
| pentamidine | SAT1 | 0.28 |
| arginine | NOS3 | 0.28 |
| macimorelin | GHSR | 0.28 |
| sildenafil | PDE1B | 0.28 |
| menadione | CDC25B | 0.28 |
| crizotinib | INPPL1 | 0.28 |
| gilteritinib | ACSL5 | 0.28 |
| ezogabine | KCNQ4 | 0.27 |
| teriparatide | PTH1R | 0.27 |
| sunitinib | PDXK | 0.27 |
| dicumarol | PCSK7 | 0.27 |
| haloprogin | TACR2 | 0.27 |
| palbociclib | CCNH | 0.26 |

| | | |
|---|---|---|
| fedratinib | CDKL5 | 0.26 |
| imatinib | KARS1 | 0.26 |
| dronabinol | GPR18 | 0.26 |
| orlistat | DAGLB | 0.26 |
| canagliflozin | SLC5A1 | 0.26 |
| zafirlukast | LTB4R | 0.26 |
| protirelin | TRHR | 0.26 |
| pyrvinium | SYNJ2 | 0.26 |
| niacin | HCAR2 | 0.26 |
| olaparib | PARP10 | 0.25 |
| omeprazole | ATP12A | 0.25 |
| warfarin | ALB | 0.25 |
| clotrimazole | CCR4 | 0.25 |
| pimozide | USP1 | 0.25 |
| miconazole | GRM6 | 0.25 |
| lindane | GABRR1 | 0.25 |

Supplementary Figure 1:

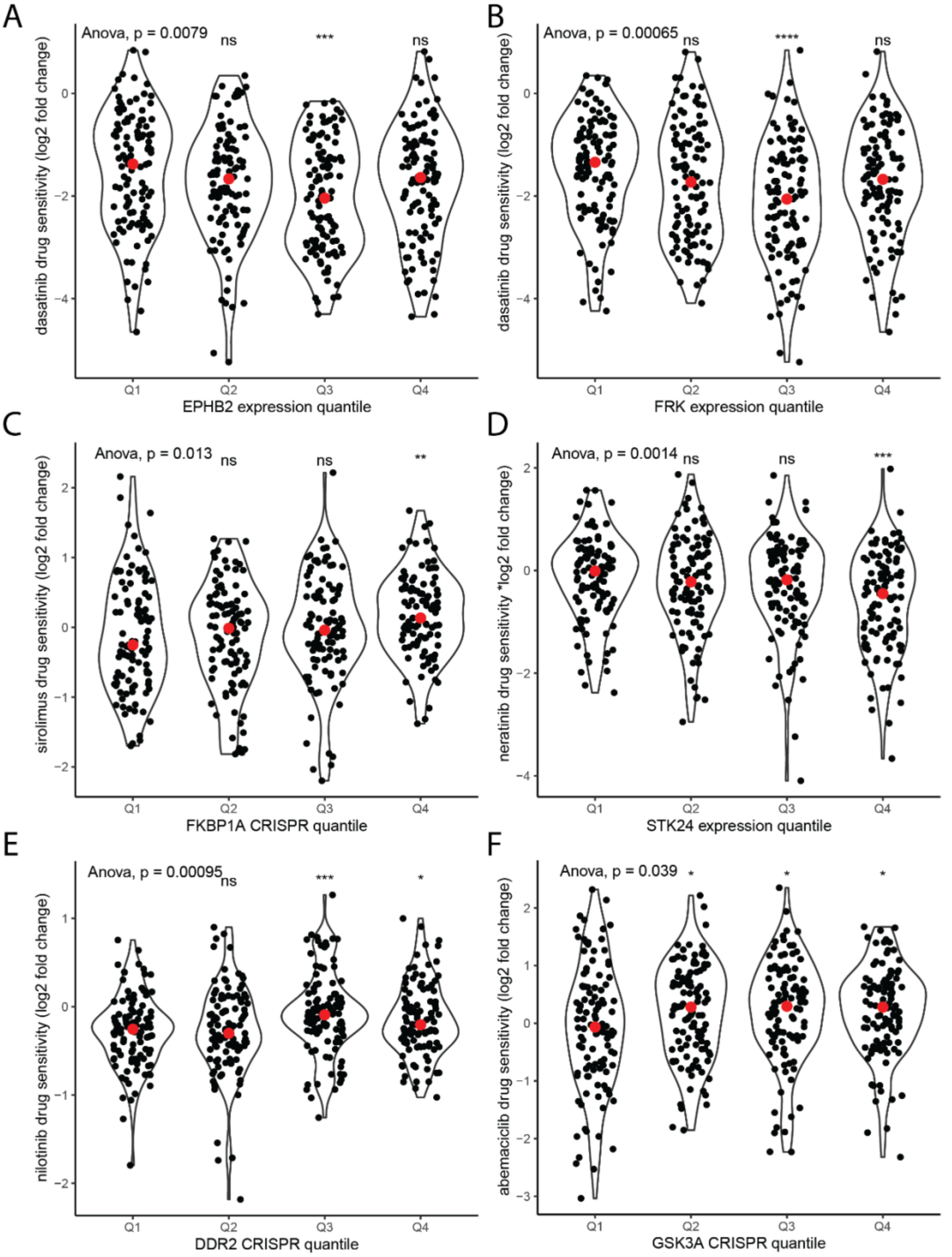

*A) Drug sensitivity for dasatinib (log2fold change in cell viability) according to EPHB2 expression B) Drug sensitivity for dasatinib (log2fold change in cell viability) according to FRK expression C) Drug sensitivity for sirolimus (log2fold change in cell viability) according to FKBP1A CRISPR gene-dependency D) Drug sensitivity for neratinib (log2fold change in cell viability) according to STK24 expression E) Drug sensitivity for nilotinib (log2fold change in cell viability) according to DDR2 CRISPR gene-dependency F) Drug sensitivity for abemaciclib (log2fold change in cell viability) according to GSK3A CRISPR gene-dependency. Asterisks indicate results of t-tests comparing relevant quartile with the first quartile of gene expression/gene dependency: * p<0.05, **p<0.01, ***p<0.001, ****P<0.0001*
